# Integrin α3β1 is required for alveolar-capillary barrier formation through regulation of epithelial differentiation

**DOI:** 10.64898/2026.10.05.755686

**Authors:** Allison D. Henry, Annica L. Alwine, Matthew J. Niederhuber, Anika K. Yarlagadda, Nikita R. Muthakana, Angie L. Mordant, Jason J. Gokey, Jennifer M.S. Sucre, Austin J. Hepperla, Jean-Philippe Cartailler, Elle N. Franciscus, Peter M. Gulleman, Alexandria L. Sharkey, Naoko Boatwright, Seunghyi Kook, Jessica C. Lee, Christopher S. Jetter, Shawyon P. Shirazi, Dawn C. Newcomb, Rachel C. Hart, J. Jeffrey Reese, Arnoud Sonnenberg, Rachel Lennon, Timothy S. Blackwell, Roy Zent, Erin J. Plosa

## Abstract

Laminin-binding integrins regulate critical morphogenic processes, but their specific roles in lung development are unknown. In this study, we evaluated a novel in vivo model of epithelial α3 subunit deletion in the murine lung with a multiomics approach to investigate the role of the epithelial α3β1 integrin during murine lung development. We demonstrate that α3β1 integrin is dispensable for airway branching morphogenesis. Instead, in vivo deletion of α3β1 integrin impaired alveolar epithelial cell differentiation and disrupted alveolar-capillary barrier formation, functionally limiting gas exchange. A discontinuous alveolar BM contained abnormal collagen IV and laminin networks and widened the gap between epithelial and endothelial cells in the alveolar-capillary barrier niche. Mechanistically, mice deficient in epithelial α3β1 integrin possessed an intracellular trafficking and secretion defect for basement membrane components. Thus, integrin α3β1 orchestrates the temporal maturation of alveolar epithelial cells required for organization of the alveolar-capillary barrier.

**Summary statement:** Integrin α3β1 is required for alveolar-capillary barrier formation through regulation of alveolar epithelial differentiation and alveolar BM composition.

## Introduction

The principal function of the lung is to exchange gas across a structurally precise alveolar-capillary barrier in the distal lung parenchyma. Unlike airway branching morphogenesis, the molecular mechanisms and the temporal regulation underpinning the creation of the alveolus remain largely undefined. Recent studies demonstrate that epithelial and endothelial cell differentiation establishes the cellular organization of the alveolar-capillary barrier niche during the early saccular stage (1–6), with organizational refinement and expansion during alveolarization (7, 8). During murine lung development, alveolar epithelial lineage specification begins at E16.5 and produces a full complement of cuboidal surfactant-producing type 2 cells (AT2) and flat gas-exchanging type 1 cells (AT1) by mid-sacculation. Notably, alveolar epithelial cells markedly change their cell shape and identity by passing through a transient cytokeratin 8+ intermediate cell state (9) before attaining the flattened cell shape of fully mature AT1 cells. In response to AT1-derived signals, general capillary endothelial cells (gCap) differentiate into alveolar capillary endothelial cells (aCap) (1, 5, 10, 11). Despite our increased understanding of the cellular components of the alveolar-capillary barrier niche, the temporal regulation and molecular mechanisms that drive maturation of the alveolar basement membrane (BM) during sacculation are entirely unknown.

Like all BMs, the alveolar BM is composed of layered laminin and collagen IV networks held together by associated BM proteins (notably nidogen and heparan sulfate proteoglycans) (12, 13). BMs serve a variety of tissue-specific functions, including filtering, sealing a barrier, stabilizing adjacent cellular compartments, or directing adjacent cellular processes. While most BMs contain the same basic components, the exact blend of laminin and collagen isoforms determine the function of the BM (14–16). Three laminin subunits, designated αβγ, combine to form laminin αβγ heterotrimer isoforms, with laminin-511 (α5β1γ1), the closely related laminin-521 (αβ52γ1) and laminin-332 (α3β3γ2) as the most abundant isoforms in the adult murine distal lung. Similarly, collagen IV is also a heterotrimer composed of three α chains (selected from α1-α6 chains). The ubiquitous collagen IV α1α1α2 network is the primary and most abundant type of collagen IV found in lung BMs, along with the less abundant collagen IV α3α4α5 network present in the alveolar BM. Epithelial cells interact with these matrix networks through integrins, heterodimeric transmembrane receptors composed of an αβ pair. Of the three laminin-binding integrins α3β1, α6β1, and α6β4, that are expressed by the developing alveolar epithelium, integrin α3β1 is the best described to direct epithelial cell behavior in organogenesis of other epithelial tissues (17, 18).

Recent reports of human variants in the *ITGA3* gene support a critical role for integrin α3β1 in lung development (19–21). Almost all the afflicted individuals die from respiratory failure and/or kidney failure in infancy or early childhood, often with notable structural abnormalities in the distal lung (19–21). Similarly, global murine deletion of the integrin α3 subunit, which uniquely targets the laminin-binding integrin α3β1 receptor, results in abnormal glomerular BM organization and dilated distal lung structures, but the alveolar-capillary barrier phenotype was not reported (22). Additional evidence that α3β1 integrin in the lung epithelium might be important for distal lung development comes from our previous murine deletion of epithelial β1 integrin, which heterodimerizes with 8 α subunits including α3. These mice developed enlarged airspaces and thickened alveolar septa suggesting impaired development of the alveolar septa (23). To date, the only murine study that investigated the role of integrin α3β1 in the lung epithelium reported normal lung development (24), leaving the role of integrin α3β1 in lung development unresolved.

In this study, we generated mice with an epithelial specific deletion of the α3 integrin subunit from E9.5. Surprisingly, these mice did not have a branching defect, however they were hypoxemic, which is suggestive of an alveolar-capillary barrier defect. The α3-deficient mice exhibited a markedly imbalanced AT2 to AT1 cell ratio, as well as a thickened, abnormal alveolar BM that failed to acquire the specialized BM components laminin-332 and collagen IV α3α4α5. Mechanistically, the α3β1-null AT1 cells exhibited defects in transcription and secretion of these matrix components. Taken together, these findings demonstrate that integrin α3β1 is required for BM formation in the alveolus.

## Results

### Integrin α3β1 regulates alveolar-capillary barrier organization and function during development

Pan-epithelial deletion of the integrin α3 subunit in the murine lung from E9.5 was achieved by crossing mice harboring Shh-Cre recombinase (Jax# 005622) (25, 26) with floxed α3 mice (27), termed α3KO mice. We confirmed epithelial-specific α3 deletion in α3KO lungs at postnatal day 3 (P3), as well as continued absence of the α3 protein at P28 by immunostaining and immunoblot, respectively (**Fig. S1A-C**). Given the established role for β1-containing integrins in early lung development (23, 28), we expected α3KO mice to exhibit a major branching defect. Surprisingly, α3KO mice were live born in the expected Mendelian ratios (**Fig. S1D**) and lived to adulthood with similar peripheral airways compared to control mice (orange arrows in **Fig. 1A**). Consistent with this finding, we observed equivalent airway branching in α3KO and control E15 fetal lung explants during fetal lung development (**Fig. S1E-F**), as well as similar gross morphology at E18 (**Fig. S1G**), just after the completion of branching morphogenesis. In contrast, histological examination at P28 revealed gross abnormalities in α3KO lungs beginning with the alveolar duct. The α3KO alveolar ducts were dilated without expansion of alveolar size or evidence of alveolar septal destruction and quantified as an increased alveolar duct volume density (**Fig. S1H**). The localized nature of this dilation suggests disrupted saccular stage lung development, as refinement of alveolar duct size and shape occurs during the early saccular stage (2). Distal to the alveolar duct, α3KO mice exhibited hypercellular, thick alveolar septa (green arrowheads, insets in Fig. 1A) compared to the thin septa observed in control mice (green arrows). By ultrastructural examination, these thick α3KO alveolar septa contained alveolar epithelial cells with mixed AT2 and AT1 cell features and varied cell shape, whereas flat AT1 cells were easily distinguishable from cuboidal AT2 cells in control lungs (**Fig. 1B**). In α3KO lungs, cuboidal epithelial cells containing the expected lamellar bodies (LB) were positioned within the alveolar septum, rather than located contiguous with the alveolar surface (Fig. 1B, denoted by AT2#). Further, the α3KO septa lacked epithelial cells possessing the characteristic thin AT1 cell shape and there were oblong epithelial cells with AT2-like microvilli (denoted by ^) or thick and elongated cells with lamellar bodies lining the α3KO septa (denoted by AT2+). Since the variation in epithelial cell shape and characteristics suggested significant impairment in differentiation, we evaluated relative AT2 and AT1 abundance. In RNAscope labeled P28 sections, lungs from P28 α3KO mice contained an overabundance of *Sftpc*+ AT2 cells and a paucity of *Hopx*+ AT1 cells compared to control lungs, resulting in an increased AT2/AT1 cell ratio (**Fig. S2A-B**). When we assessed cell proliferation using *Mki67* in our analysis, there was only an incremental increase in proliferating AT2 cells in α3KO lungs, and no difference at P3 (**Fig. S2C**). This finding suggested ongoing failed epithelial differentiation as the primary mechanisms for AT2 overabundance rather than unchecked AT2 proliferation. To assess the degree of alveolar epithelial differentiation achieved in the absence of α3β1 integrin, we immunostained P28 lung sections for AT2, intermediate, and AT1 cell markers. The majority of ABCA3+ AT2 epithelial cells in α3KO lungs co-expressed the intermediate cell marker cytokeratin 8 (Krt8) (arrowheads in **Fig. 1C**), while control lungs contained fewer Krt8+/ ABCA3+ dual positive epithelial cells (**Fig. S2D**). Since changing cell shape is a necessary step in alveolar epithelial differentiation, we calculated the roundness score for the ABCA3+ cells (29) and found that the α3KO ABCA3+ AT2 cells were significantly less round compared to control ABCA3+ AT2 cells (**Fig. S2E**). These data are further evidence of impaired epithelial differentiation and are consistent with the varied AT2 cell shape noted by our ultrastructural studies. We next assessed the degree of AT1 elongation and maturation by Krt8 and T1α immunostaining. Control alveolar septa contained numerous elongated T1α+ AT1 cells, occasionally co-expressing both T1α and Krt8 (**Fig. 1D**, arrows). In contrast, elongated cells in α3KO distal lungs only expressed Krt8+ (Fig. 1D, arrowheads), with a minority expressing dual T1α+/ Krt8+ markers and very rare cells expressing T1α alone, suggesting blocked intermediate to AT1 differentiation. Taken together, these findings demonstrate that α3β1 integrin regulates both AT2 to intermediate and intermediate to AT1 differentiation.

**Figure 1.**
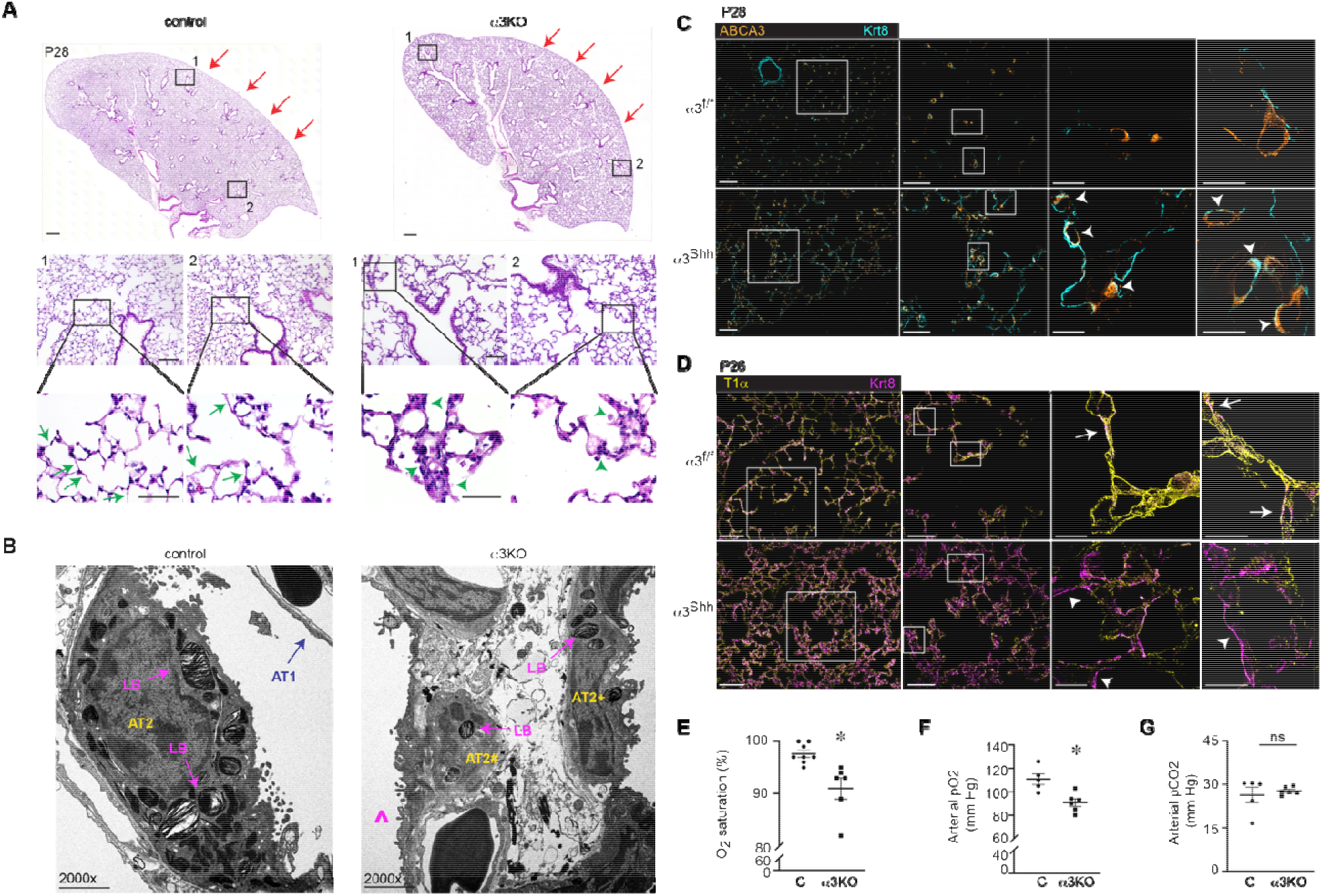
Epithelial α3β1 integrin is required for epithelial differentiation and function of the alveolar-capillary barrier. (A) Structural deficits in α3KO lungs are observed in the distal lung parenchyma. Stitched whole lobe 20x images demonstrate similar size and orientation of peripheral airways between the genotypes (orange arrows). High-power insets show thickened alveolar septa (green arrowheads) in α3KO lungs and typical thin alveolar septa in control lungs (green arrows). (B) P28 TEM images show distinct cuboidal AT2 and flat AT1 cells in control lungs. α3KO lungs contain alveolar epithelial cells with varied cell shape and mixed AT1/AT2 features (LB= lamellar bodies, ^ flat epithelial cell with microvilli, AT2+ thick elongated AT2-like cell with LB and microvilli, AT2# intraseptal cuboidal AT2-like cell). (C) Representative images of P28 lung sections immunostained for Krt8 (cyan) and ABCA3 (orange) demonstrating increased dual ABCA3+/ Krt8+ epithelial cells in α3KO distal lung, compared to control lungs. (D) Representative images of P28 lung sections immunostained for Krt8 (magenta) and T1α (yellow) showing increased mature T1α+ AT1 cells in control lungs and increased Krt8+ epithelial cells in α3KO lungs. Krt8/ T1α co-localization marked by arrows, numerous Krt.8+ epithelial cells in α3KO lung marked by arrowheads. (E) Decreased O2 saturation in 3-month-old α3KO mice by MouseOx measurement, 91.0±2.0% in α3KO vs. 97.6±1.8% in control (n = 8 control, 6 α3KO mice). (F-G) iStat arterial blood gas analysis shows decreased paO2 (90.8±3.2 mmHg for α3KO vs. 110.8±4.6 mmHg for control) and equivalent paCO2 (27.8±0.5 mmHg for α3KO vs. 26.4±2.7 mmHg for control) in conscious 3-month-old α3KO mice; n = 5 control, 6 α3KO mice. For A, scale bar = 500 μm for stitched whole lobe 20x images; 100 μm for low-power images; 50 μm for high power insets. For B, scale bar = 2 μm for 2000x; direct magnification as indicated. For C and D, scale bar = 100 μm for stitched whole lobe 20x images. 50 μm for low power, 10 μm for insets. * *p* < 0.05 by two-tailed Welch’s t-test.

Since the capacity to exchange gas depends on the structure and cellular organization of the alveolar septum, we next tested for physiologic impairment specifically attributable to alveolar-capillary barrier dysfunction. We detected mild hypoxemia by MouseOx measurement in α3KO mice at 3 months of age (**Fig. 1E**), and they exhibited decreased compliance and increased resistance on FlexiVent lung function testing (**Fig. S2F**), suggesting restrictive functional impairment. We next more precisely tested for differences in oxygen diffusion through arterial blood gas analysis. Arterial samples were drawn from conscious 3-month-old mice via indwelling femoral arterial catheters, thereby avoiding supplemental oxygen administration that could obscure detection of diffusion. As anticipated given the alveolar septal thickness, we detected marked hypoxemia in α3KO mice compared to control mice (α3KO paO_2_ 90.8±3.2 mmHg vs. 110.8±4.6 mmHg paO_2_ for control mice; **Fig 1F**). Importantly, we observed equivalent paCO_2_ between the genotypes (**Fig. 1G**), which suggests impaired diffusion across the alveolar BM in α3KO mice. Taken together, these data demonstrate that epithelial α3β1 is required for proper epithelial differentiation, cellular organization, and function of the alveolar-capillary barrier.

### Integrin α3β1 is required for alveolar BM formation during sacculation

Clear deficits in alveolar-capillary barrier organization and function suggested that α3β1 integrin regulates formation of the alveolar BM, the specialized matrix structure at the center of the alveolar-capillary barrier niche. We examined this niche by capturing serial transmission electron microscopy (TEM) images during distal lung development beginning at E18. We measured the width of the alveolar BM between AT1 and capillary endothelial cells, standardized at the level of the alveolar duct opening (**Fig. 2A**). At E18 and P0, a gap existed in the BM space between epithelial and endothelial cells for both genotypes (**Fig. S3A**). However, the alveolar BM in control lungs thinned by P3, remaining consistently thinner than the α3KO alveolar BM during alveolarization at P5 and P28 (**Fig. 2B-C, Fig. S3B**). Notably, the failure to narrow the alveolar BM space by P3 in α3KO lungs was followed by progressive enlargement from P5 to P28. The P3 nadir also occurred in control lungs, albeit more subtle, suggesting the alveolar BM is remodeled during alveolarization in normal lung development. In summary, these data precisely define both when alveolar BM formation occurs in WT murine lungs and identifies P3 as a point of divergence between control and α3KO murine lungs.

**Figure 2.**
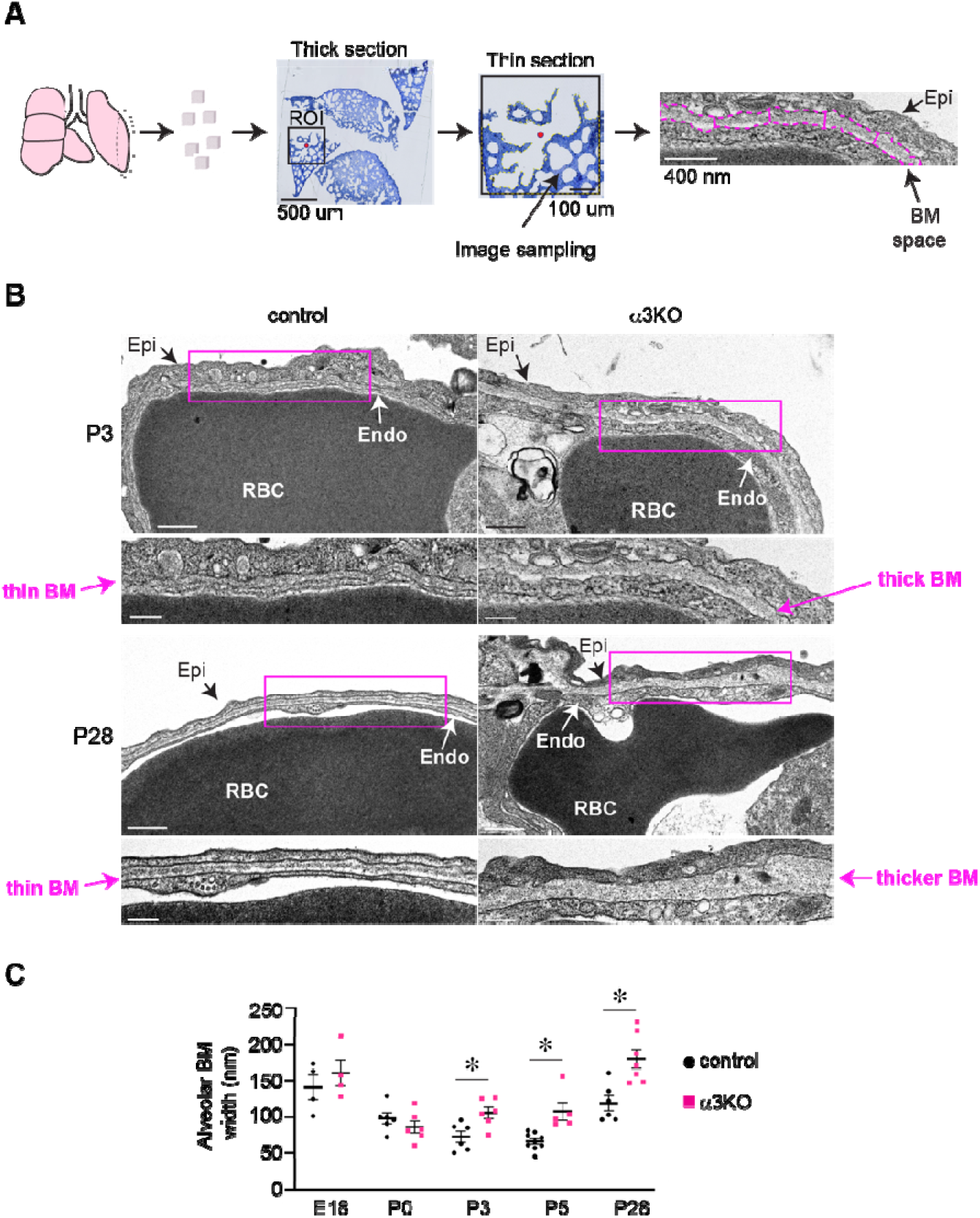
Integrin α3β1 is required for alveolar BM formation during sacculation. (A) Schematic of standardized alveolar BM measurements by TEM. After thick sectioning, selection of a region of interest (ROI) centered at the alveolar duct entrance (dot) was thin sectioned, images were sampling from the adjacent post-ductal peripheral lung. The width of the alveolar BM space was quantified by perpendicular measurements every 400 nm along the epithelial-endothelial alignment, defined by an associated intra-capillary red blood cell (RBC). (B) TEM developmental time series demonstrates acquisition of a thin alveolar BM by P3 in control mice. Disruption of alveolar BM formation in α3KO lungs is observed by P3, resulting in progressive enlargement of the BM space. Additional images for E18, P0, and P5 timepoints in Fig. S3A, B. (C) Measurement of alveolar BM width (≥ 8 interval measurements per 5000x magnification section, sampled 400 nm apart, n = 4-9 mice/ genotype at each timepoint as indicated by plot and each dot representing the mean width per individual mouse, 5-6 sections/ mouse). Genotypes were compared by two-tailed Welch’s t-test at each timepoint, * *p* < 0.05. Scale bar = 500 nm for low-power TEM (top panels in B), 200 nm for TEM insets (bottom panels in B).

Having defined when α3β1 regulates alveolar-capillary barrier formation, we next examined *Itga3* expression levels in developing wild-type (WT) murine lungs to identify the epithelial cell population in which α3β1 integrin is required for distal lung development (**Fig. 3A**). Querying our published developmental scRNA-sequencing atlas, we noted robust *Itga3* expression in epithelial and endothelial populations, with negligible expression in fibroblasts in WT murine lung throughout development. Among the epithelial cell populations, we observed modest *Itga3* expression in the pre-saccular lung, primarily restricted to early AT2 cells. However, *Itga3* expression robustly increased at the start of sacculation (E18) in AT1 and intermediate cells. We confirmed α3 subunit protein localization in AT1 cells postnatally in WT lungs by immunostaining (**Fig. 3B**, single channels in **Fig. S3C**). During early sacculation (P0), both AT2 (marked by ABCA3, yellow arrows) and early AT1 cells (marked by T1α, white arrows) expressed the α3 protein (magenta). During alveolarization (P7 and P14), α3 primarily localized to AT1 cells, validating the WT transcriptomic data. Taken together, the timing and localization of *Itga3* expression in the developing WT lung suggested that α3β1 integrin has a specialized role in AT1 cells from mid-sacculation and into alveolarization.

**Figure 3.**
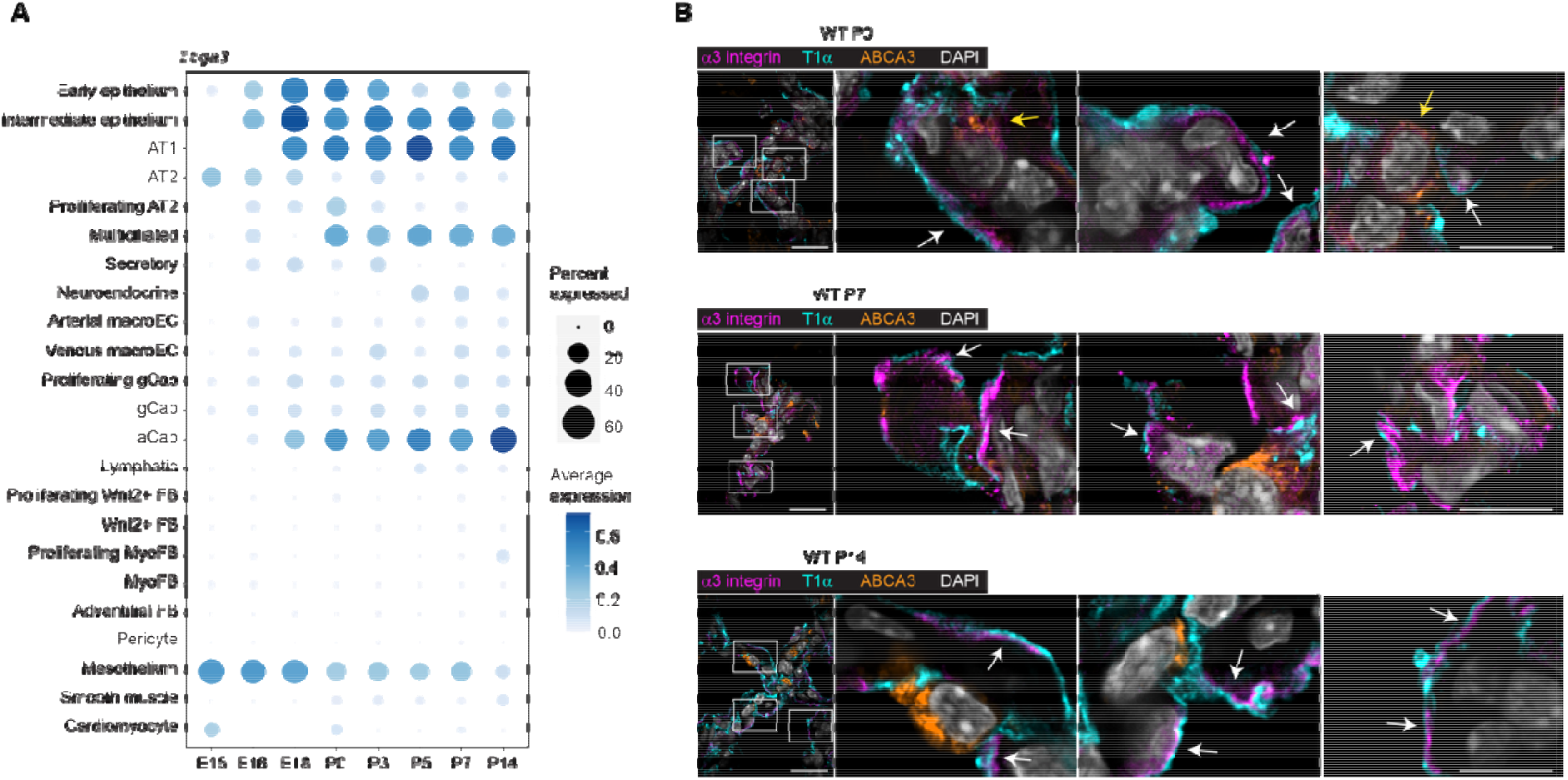
WT *Itga3* expression abruptly increases during sacculation and shifts to AT1-predominance during alveolarization. (A) Dot plot representing *Itga3* expression from WT murine lung demonstrating increased AT1 and intermediate cell expression of *Itga3* during sacculation and alveolarization. (B) Lung sections immunostained for the α3 integrin subunit (magenta), T1α (cyan), and ABCA3 (orange) shows co-localization of α3 integrin with both AT2 and AT1 cells during early sacculation (P0) but primarily with AT1 cells during alveolarization (P7 and P14). Single channel images in Fig. S3C. Scale bar = 25 μm for low-power, 5 μm for high-power insets in B.

### Integrin α3β1 regulates saccular stage alveolar epithelial differentiation

We next assessed α3KO and control lungs for alveolar epithelial differentiation defects at P3, the point at which α3β1 integrin critically regulates alveolar BM formation. Consistent with P28 histology, P3 α3KO lungs contained thickened interstitial septa when compared to control P3 lungs. (**Fig. 4A**, quantified by mean interstitial intercept in **Fig. S4A**). We did not detect a difference in the ratio of *Sftpc+/ Hopx+* cells or the percentage of proliferating AT2 cells between the genotypes at P3 (**Fig. 4B**, **Fig. S4B-C**). However, a higher percentage of *Sftpc*+ AT2 cells clustered in pairs or triplets were seen within α3KO alveolar septa when compared to control alveolar septa (arrows in Fig. 4B, quantified in **Fig. S4D**). This atypical arrangement suggests that α3β1 integrin is required for epithelial migration, which was confirmed by in vitro migration assays on laminin-511 and -332, the BM components normally positioned closest to the lung epithelial layer (**Fig. S4E-F**).

**Figure 4.**
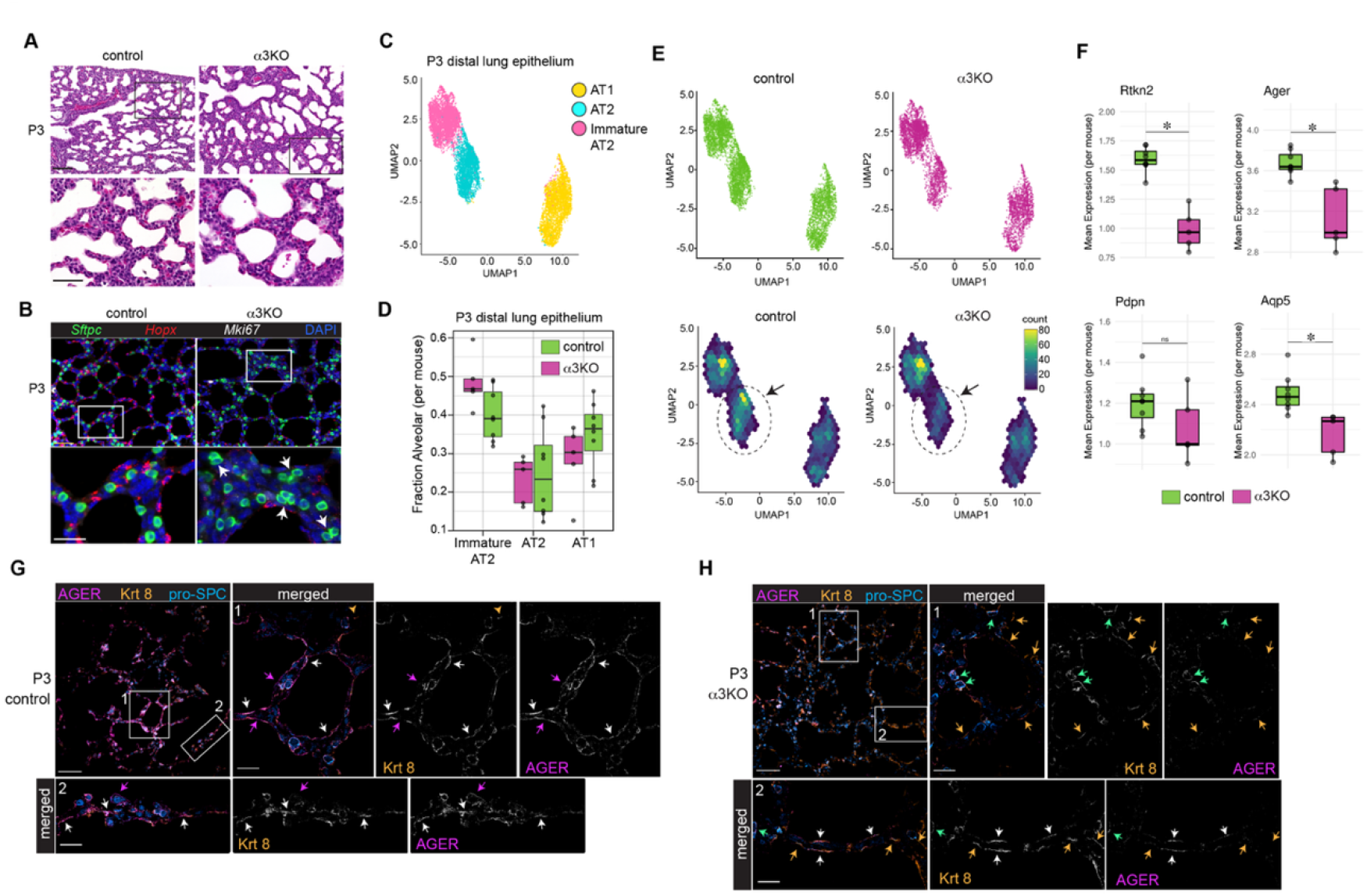
Integrin α3β1 regulates alveolar epithelial organization and differentiation during sacculation. (A) H&E-stained lung sections demonstrate thickened alveolar septa in P3 α3KO mice. (B) *Sftpc*+ AT2 cells cluster in pairs and triplets in P3 α3KO lungs (arrows) but disperse as single cells in P3 control lungs. Cells identified by RNAscope probes for *Sftpc, Hopx,* and *Mki67.* (C) Unified Manifold Approximation and Projection (UMAP) plot of single cell transcriptomic profiles showing 3 distinct alveolar epithelial cell clusters at P3. (D) Box plots comparing per-mouse alveolar subtype fractions (black points) relative to all alveolar cells reveals a trend towards increased *S100g*+ immature AT2 cells in α3KO lungs. (E) UMAP plots split by genotype and labeled by hexagonal bins of cell density. Arrows and dashed circles highlight decreased mature AT2 cell density in P3 α3KO lungs. (F) Box plots with per-mouse average (mean) single-cell gene expression (SCT normalized) show decreased expression of AT1 hallmark genes *Rtkn2, Ager,* and *Aqp5* in P3 α3KO mice. For C-F, n = 5 control mice, 8 α3KO mice. (G-H) Representative images of lung sections immunostained for pro-SP-C (blue), Krt8 (orange), and AGER (magenta). Elongated AGER+ AT1 cells are observed in control lungs (purple arrows), as well as elongated dual AGER+/Krt8+ epithelial cells (white arrows). Lungs from α3KO mice contain AGER-/Krt8+ epithelial cells (orange arrows), dual AGER+/ Krt8+ cells, and cuboidal triple AGER+/ Krt8+/ pro-SP-C+/ epithelial cells (green arrows), but AGER+/ Krt8-cells are absent in α3KO lungs. Scale bar for A= 100 μm for low-power, 50 μm for inset. Scale bar for B= 50 μm for low-power, 25 μm for inset. Scale bar for G and H = 100 μm for low-power, 20 μm for inset 1 and 2. * *p* ≤ 0.05 by Welch’s t-test.

As *Hopx* might lack specificity as an AT1 marker at P3, we examined AT2, intermediate cell, and AT1 transcriptional phenotypes by scRNA-sequencing analysis. After hierarchical lineage assignment by established hallmark genes, we identified the primary *Sftpc+* AT2 and *Rtkn2+* AT1 alveolar populations. Subsequent analysis of markers that defined an unbiased clustering of these cells identified a third alveolar subset primarily marked by expression of *S100g* and *Sftpc* (**Fig. S4G-I**). *S100g is* a vitamin D pathway gene expressed at E19-20 in rat alveolar epithelial cells, just prior to saccular stage birth (30), suggesting a block in AT2 maturation. We classified these *S100g+* AT2 cells as an “immature AT2” subset (**Fig. 4C**). Quantification of alveolar epithelial fractions across all three subtypes (immature AT2, AT2, AT1) demonstrated a 1.2x increase in the median fraction of immature AT2 cells (p-value = 0.067, Welch’s t-test) and a 0.8x decrease in the median fraction of AT1 cells (p-value = 0.24, Welch’s t-test) in P3 α3KO mice relative to controls (**Fig. 4D**). Consistent with this finding, UMAPs of cluster density revealed a paucity of fully differentiated AT2 cells in P3 α3KO lungs (dashed circle and arrow in **Fig. 4E**). Expression of the AT1 markers *Rtkn2*, *Ager,* and *Aqp5* were significantly decreased in the AT1 population of α3KO versus control lungs (*p*-value ≤ 0.05, Welch’s t-test; **Fig. 4F**), providing further transcriptional evidence of impaired alveolar epithelial differentiation with loss of α3β1 integrin.

We next examined P3 epithelial cell shape and validated our scRNA-seq analysis by immunostaining for pro-SP-C (blue), Krt8 (orange), and AGER (magenta), respectively. In control lungs, elongated AGER+Krt8-epithelial cells lined the alveolar septa (purple arrows), indicative of the presence of mature AT1 cells (**Fig. 4G**). These cells were mixed with flat dual AGER+/Krt8+ (white arrows) representing less mature intermediate-AT1 cells. In contrast, α3KO lungs contained numerous elongated AGER-/Krt8+ cells (orange arrows in **Fig. 4H**), occasional elongated AGER+/ Krt8+ cells (white arrows), and rare cuboidal triple positive AGER+/Krt8+/ pro-SP-C+ cells (green arrows). However, there was a notable absence of only AGER+ cells in α3KO lungs, consistent with impaired AT1 maturation. Similarly, we confirmed continued epithelial ambiguity during alveolarization by immunostaining for the AT1 marker T1α with pro-SP-C (**Fig. S4J**). Dual T1α+/ pro-SP-C+ epithelial cells were observed at P14, well past the expected resolution of primordial epithelium (31–33). Taken together, these histological and transcriptional data indicate that integrin α3β1 is required for the organization and differentiation of alveolar epithelial cells during sacculation.

### Epithelial α3β1 integrin directs alveolar BM composition

Since AT1 cells serve as the exclusive source of certain collagen IV and laminin isoforms found in the alveolar BM (34) and α3KO lungs contain fewer mature AT1 cells, we hypothesized that mice lacking integrin α3β1 would have impaired alveolar BM formation due to decreased secretion of BM matrix components. We first examined the localization of collagen IV α1α1α2 and laminin-511/521 isoforms, which are expressed by multiple cell types, in the alveolar-capillary barrier niche. As expected, these subunits co-localized neatly with AGER+ epithelial cells in control lungs (white arrows for white overlay) (**Fig. 5A**). In contrast, α3KO lungs exhibited a marked increase in these isoforms within the alveolar septa and intracellularly in ABCA3+ AT2 cells, with very few areas of co-localization with AGER+ epithelial cells. As we perceived a difference in protein abundance, we next determined if loss of epithelial integrin α3β1 changed transcription of the subunits comprising these isoforms. We analyzed a scRNA-seq dataset from adult control and α3KO lungs using a combined AT1/ intermediate cluster due to small AT1 yield (UMAPs of clusters and epithelial cell annotations defined in **Fig. S5A-B**). All three alveolar-capillary barrier niche cell types expressed collagen IV α1α1α2 and laminin-511/521 genes, and, consistent with the immunostained lungs, α3KO aCap EC and AT1/ intermediate cells exhibited increased expression of the majority of the subunits (**Fig. 5B**). We next examined localization and expression of the subunits comprising collagen IV α3α4α5 and laminin-332, isoforms which have been associated with thinning of the glomerular BM (known as isoform switching) (16, 35–37). In adult control lungs, the collagen IV α3 and laminin β3 subunits tightly co-localized with AGER+ AT1 cells (arrows in **Fig. 5C**), however, in α3KO lungs, both laminin β3 and collagen IV α3 localized intracellularly to cuboidal ABCA3+ epithelial cells, with minimal deposition along AGER+ epithelium (arrowheads). Similarly, in control lungs laminin γ2 (red) was present along the thin alveolar BM that was identified by nidogen-1, a critical BM assembly component (**Fig. 5D**, arrowheads, left panel). In contrast, α3KO ABCA3+ epithelial cells retained laminin γ2 intracellularly (arrows, right panel) and the cells were interspersed amongst markedly increased (and suspected compensatory) nidogen-1 deposition. We next used our adult scRNA-seq data set to address which cell types within the immediate alveolar BM niche express collagen IV α3α4α5 and laminin-332 subunit genes in the context of integrin α3β1 deficiency (**Fig. 5E**). Consistent with our prior WT data during lung development, expression of *Lama3, Lamb3*, *Lamc2, Col4a3*, *Col4a4*, and *Col4a5* was universally absent from aCap ECs in both genotypes (34). Since expression of these isoform genes remained restricted to AT1/ intermediate cells in α3KO lungs, we concluded that AT1-lineage epithelial cells serve as the exclusive and requisite cellular source for these matrix components and there was no compensation by neighboring cells. The transcriptional specificity, intracellular localization of collagen IV α3α4α5 and laminin-332 subunits and their notable absence from the alveolar septum in α3KO lungs suggest that integrin α3β1 expression in alveolar epithelial cells is required for acquisition of the proper components of the alveolar BM.

**Figure 5.**
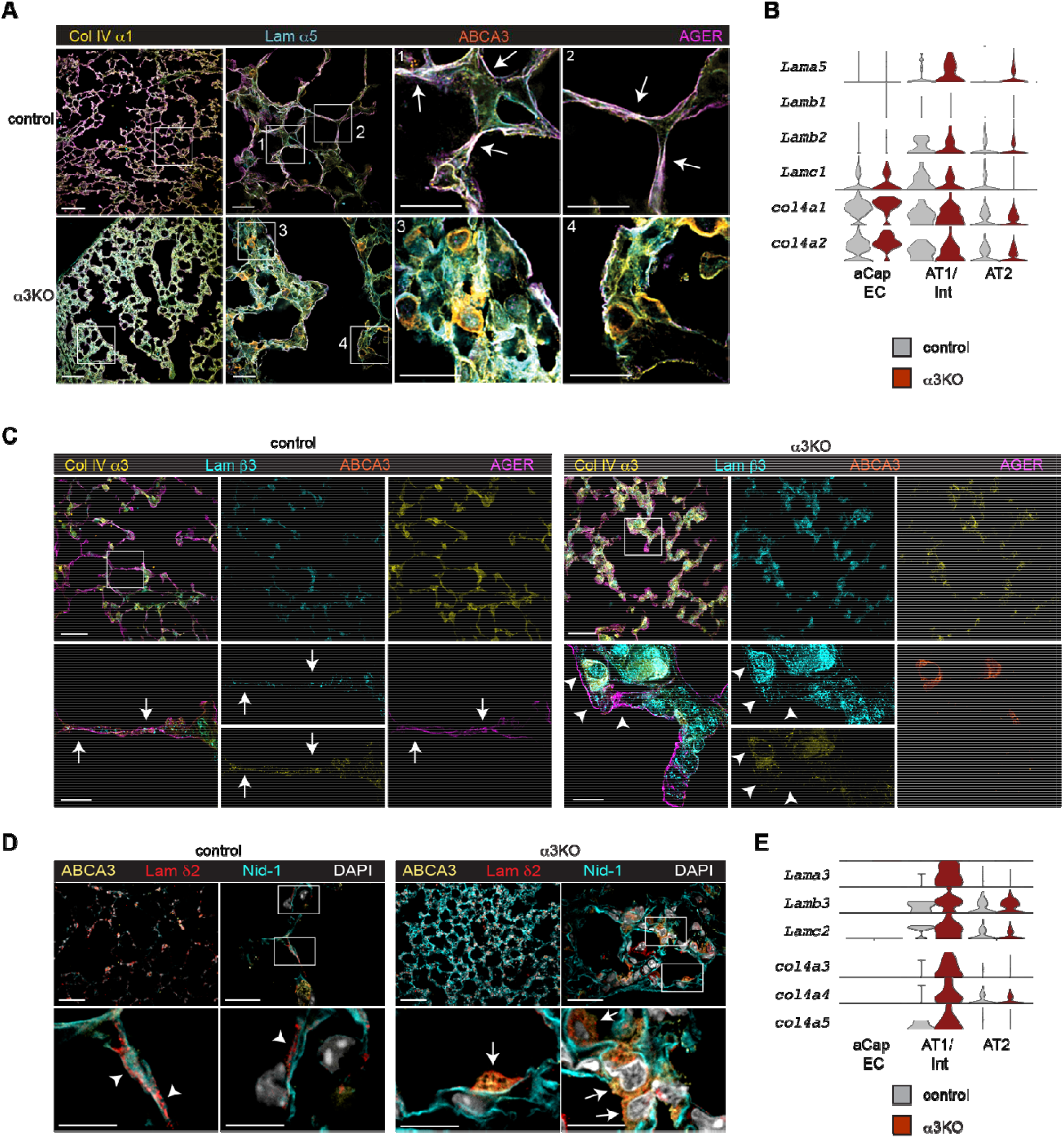
Epithelial α3β1 integrin regulates alveolar BM composition. (A) Representative immunostaining for collagen IV α1 (yellow) and laminin α5 (cyan) shows tight co-localization with AGER+ (magenta) AT1 cells in P28 control lungs (white arrows). P28 α3KO lungs contain excess collagen IV α1 and laminin α5 within the alveolar septa and ABCA3+ AT2 cells (orange). (B) Violin plots demonstrate collagen IV α1α1α2 and laminin-511/521 subunit expression in multiple cell types in the alveolar-capillary barrier niche in 3-month-old adult control and α3KO mice. n= 4 control and 3 α3KO mice. (C) In representative P28 immunostained lung sections, collagen IV α3 and laminin β3 subunits co-localize with AGER+ AT2 cells along the alveolar septum in control lungs (white arrows). In P28 α3KO lungs, collagen IV α3 and laminin β3 collects intracellularly in ABCA3+ epithelial cells, remote from elongated AGER+ cells (arrowheads). (D) White arrowheads show laminin γ2 (red) deposited along the length of control alveolar septa (identified by BM assembly protein Nidogen-1 in cyan). In α3KO lungs, laminin γ2 co-localizes intracellularly with ABCA3+ α3KO epithelial cells (white arrows) but is largely absent from the length of α3KO alveolar septa. (E) Violin plots show collagen IV α3α4α5 and laminin-332 subunits are exclusively expressed by both control and α3KO AT1/ intermediate epithelial cells but are absent from endothelial cells from both genotypes in the alveolar-capillary barrier niche. n= 4 control and 3 α3KO mice. Scale bar for A = 100 μm for low power images; 10 μm for high-power images; 5 μm for alveolar septal insets 1-4. Scale bar for C = 50 μm for low power (top row), 10 μm for insets (bottom row). Scale bar for D = 100 μm for low power, 25 μm for high power, and 10 μm for insets (bottom row).

To confirm that integrin α3β1 expression by alveolar epithelial cells is required for regulating secretion of laminin-332 and collagen IV α3α4α5 at P3, we examined the relative abundance of collagen IV α3α4α5 and laminin-332 proteins by label-free mass spectrometry-based data-independent acquisition (DIA) proteomics, a technique well suited for detection of low abundant proteins. We fractionated P3 whole lung by solubility (38), generating an extracellular fraction enriched in BM proteins and a cellular fraction composed of intracellular and transmembrane proteins. Principal component analysis (PCA) demonstrated that the fractions primarily cluster by genotype, with the α3KO cellular proteome distinct from the control proteome at P3 (**Fig. S6A**). We identified 10,840 unique proteins in total, of which 10,267 were non-matrisome proteins, 80 were considered contaminants, and 493 were categorized as lung matrisome proteins, as defined by Naba et al., 2012 (39) (**Fig. 6A**). Of the matrisome proteins identified, approximately half were categorized as ECM regulators or glycoproteins (**Fig. 6B**). The remaining matrisome proteins included collagens, proteoglycans, secreted factors, and ECM affiliated proteins. We detected all but one of the individual subunit chains (collagen IV α5) that comprise laminin-332 and collagen IV α3α4α5. We suspect this omission is due to the status of collagen IV α5 as an unverified sequence in the UniProt database at the time of proteomic analysis. We calculated relative abundance by the ratio of log_2_ fold change in α3KO to control lungs (**Fig. 6C**), demonstrating increased laminin-332 and collagen IV α3α4α5 chains in the cellular fraction of P3 α3KO lungs, with all three laminin chains reaching significance. Conversely, the extracellular fraction contained less collagen IV α3α4α5, as marked by significantly decreased collagen IV α3 and α4 abundance (**Fig. 6C**, additional visualization by volcano plot in **Fig. S6B**). Together, the paucity of collagen IV α3 and α4 in the α3KO extracellular space, combined with the cellular accumulation of laminin-332, suggests two possibilities. One explanation is increased protein synthesis of BM components, and another possibility is that integrin α3β1 has a role in regulating matrix secretion. To observe patterns of systemic dysregulation, we leveraged the Kyoto Encyclopedia of Genes and Genomes (KEGG) database of curated functional pathways to map proteins with significant α3KO versus control abundances (q-value ≤ 0.05 in either extracellular or cellular fractions) to genes annotated as part of the Focal Adhesion pathway (KEGG mmu04510, n=202 genes). Grouping these genes by known and predicted physical relationships estimated by the STRING database, further demonstrated marked dysregulation of laminin-332 abundance, compared to other laminin isoforms, in this whole lung sample (**Fig. 6D**). In addition, we observed notable deficits in cytoskeletal networks and secretion pathways, suggestive of impaired intracellular trafficking. Hierarchical and k-means clustering of these genes, again using the proteomic log2 fold change values between α3KO and control genotypes, revealed broad dysregulation of matrix-integrin-cytoskeletal interactions, further suggesting integrin α3β1 regulates cytoskeletal-dependent intracellular trafficking in epithelial cells (**Fig. 6E**). Given substantial evidence of impaired BM component secretion with the possibility of an intracellular trafficking defect, we next investigated whether α3β1 integrin regulates vesicle-mediated secretion of BM components.

**Figure 6.**
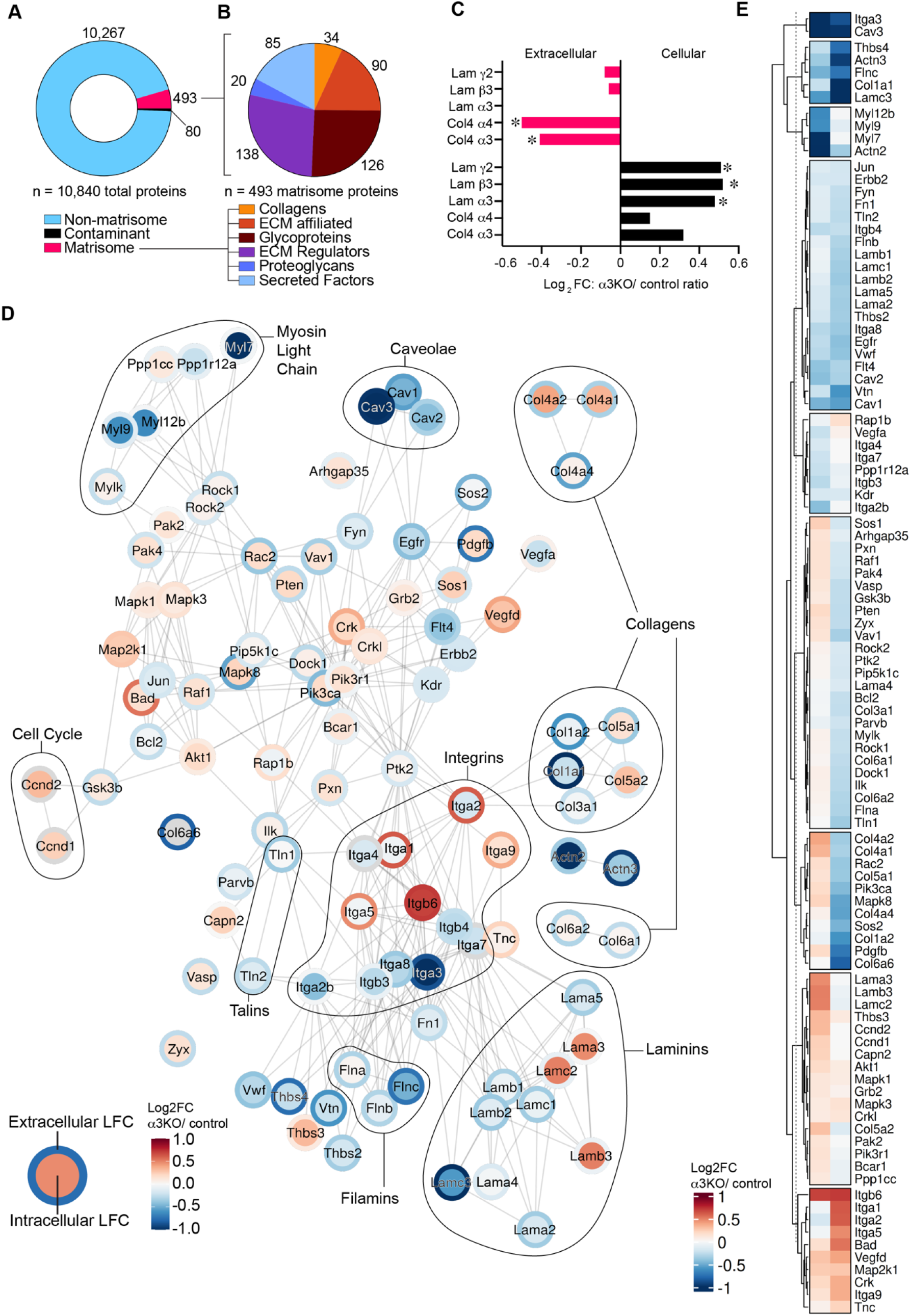
Integrin α3β1 is required for laminin-332 and collagen IV α3α4α5 secretion during sacculation. (A) Distribution of proteins identified by data-independent acquisition (DIA) proteomics in P3 control and α3KO lungs. (B) Categorization of matrisome proteins according to the Naba et al. classification (39). (C) P3 protein relative abundance calculated as log_2_ fold change of the relative abundance ratio of α3KO over control (log_2_ FC: α3KO/ control). Identification of significantly increased laminin-332 subunits in the α3KO cellular fraction and significantly decreased abundance of collagen α3 and α4 in the α3KO extracellular fraction. (D) String network of log_2_ FC: α3KO/ control comparing relative abundance in the cellular fraction (node fill color indicates cellular genotype log_2_FC) versus extracellular fractions (outer ring color indicates extracellular genotype log_2_FC). Node positioning indicates functional relationships with lines representing known or predicted protein-protein relationships (E) Hierarchical and k-means clustered heatmap of extracellular and cellular log_2_FC: α3KO/ control values in significantly differential proteins (q-value ≤ 0.05 in at least one fraction) linked to KEGG Focal Adhesion annotated genes. Relative abundance depicted as log_2_ FC: α3KO/ control from the cellular and extracellular fractions. * q-value ≤ 0.05.

### Integrin α3β1 supports polarized secretion towards the AT1 basal cell membrane

To broadly evaluate the α3β1-dependent AT1 cell functions, we examined the top 10 differentially expressed genes from our adult transcriptional data set. Consistent with the P3 proteomics data, four of the top 10 genes (*Cpm*, *Rai14*, *Ctsc*, and *Rgcc*) encode proteins involved in vesicular transport, the primary mechanism utilized for matrix protein secretion (**Fig. 7A**). Collagen IV secretion depends on vesicular size modifications by the COPII complex which can accommodate large matrix proteins (reviewed in (40)). Importantly, our analysis identified decreased expression of two COPII component genes (*Sar1a* and *Sec31a*) in α3KO AT1/ intermediate cells (**Fig. 7B**), supporting a role for AT1 α3β1 integrin in vesicle-mediated secretion of BM components. In agreement with the transcriptional data, immunostaining for Rai14, a nascent vesicle marker, revealed abundant new vesicles adjacent to AGER+ AT1 cells (**Fig. 7C**, blue arrows in insets) in control P28 alveolar septa, consistent with successful matrix protein secretion into the BM space in control lungs. In α3KO lungs, Rai14 was largely absent from the few elongated AGER+ epithelial cells (blue arrowheads in inset), but instead it was primarily localized to neighboring cell membranes (yellow arrows in insets). We next directly visualized vesicles within the epithelium by TEM and quantified the total number of vesicles per epithelial area (**Fig. 7D-E**). While we observed numerous endothelial vesicles for both genotypes (blue arrowheads), the α3KO epithelium contained fewer vesicles than control AT1 cells (magenta arrows). These findings are consistent with the concept of α3β1 integrin-dependent vesicle-mediated transport in the alveolar-capillary barrier. Since vesicles track along microtubules intracellularly and microtubules anchor to α3β1 integrin at the basal membrane through the linker protein LL5B (41), we examined localization of these microtubule-mediated transport complexes. As expected, LL5B localized along AT1 and AT2 cell membranes throughout control distal lung (white arrows), however α3KO lungs were largely devoid of LL5B, including along rare AGER+ epithelial cells (white arrowheads) (**Fig. 7F**), as quantified by fluorescence intensity per AGER ROI area (**Fig. 7G**). Thus, α3β1 integrin is required for secretion of BM components towards the basal membrane of AT1 cells.

**Figure 7.**
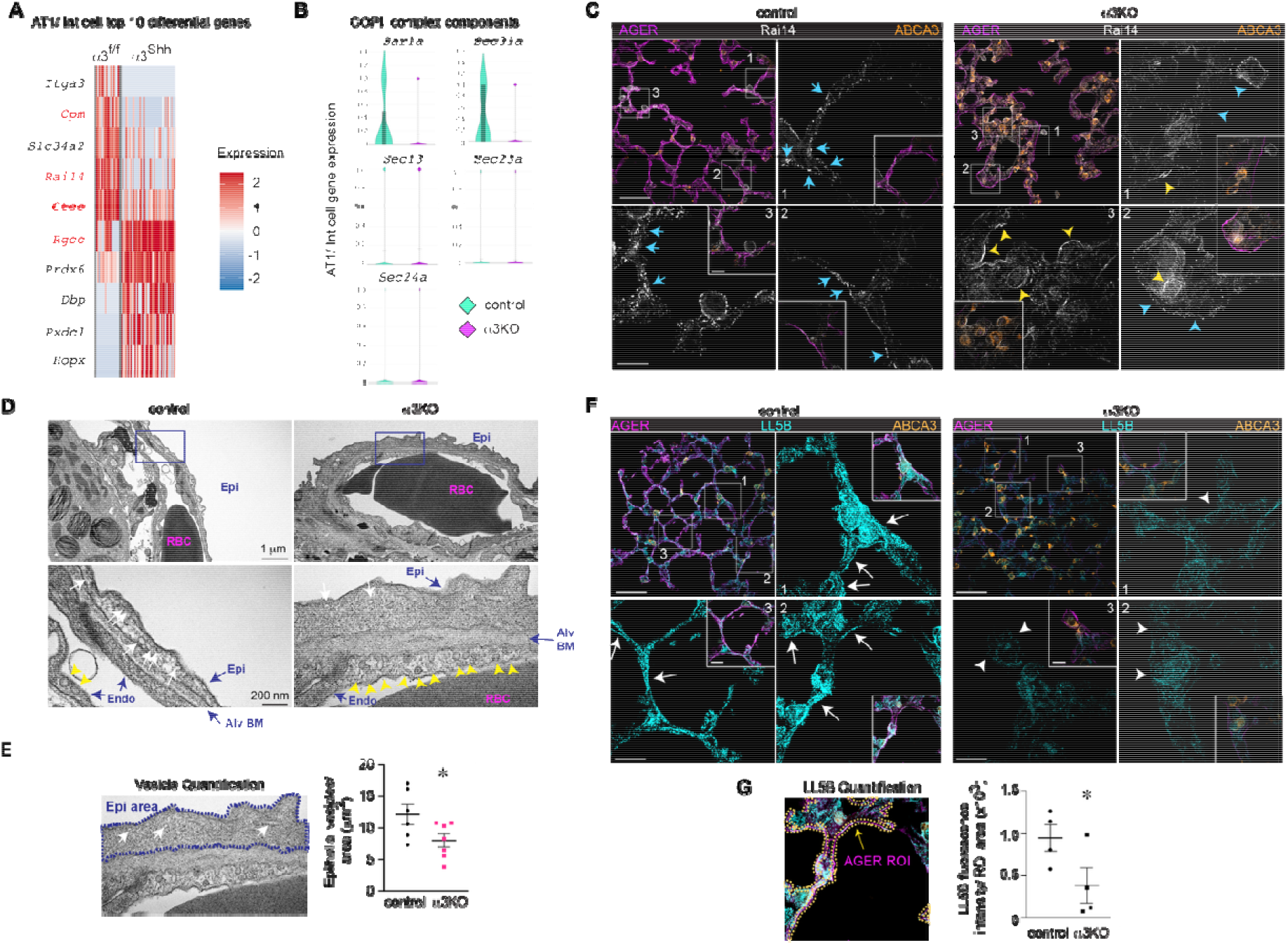
Integrin α3β1 integrin regulates vesicular transport in AT1 cells. (A) Heat map of the top 10 differentially regulated genes in adult AT1/ intermediate cells, notable for decreased expression of vesicular transport genes (*Cpm*, *Rai14*, *Ctsc,* and *Rgcc*) in α3KO AT1/ intermediate cells. (B) Violin plots demonstrate decreased α3KO AT1/ intermediate cell expression of *Sar1a* and *Sec31a*, COPII complex components required for collagen IV secretion. (C) Nascent vesicle marker Rai14 localizes to AGER+ AT1 cells in control lungs (blue arrows), while Rai14 is rarely seen along AGER+ epithelial cells in α3KO lungs (blue arrowheads). Occasional Rai14 is localized to AGER-/ ABCA3+ cells in α3KO lungs (yellow arrowheads). (D-E) Marked decrease in vesicle abundance per epithelial area in α3KO distal lungs compared to control lung, visualized by TEM. Vesicles marked by magenta arrows in epithelial cells and blue arrows in endothelial cells; n = 6 control and 7 α3KO mice. (F-G) LL5B, a protein that docks vesicle transport microtubules at α3β1 integrin on the basal membrane during exocytosis, is largely absent from in α3KO alveolar septa; quantified in AGER-adjacent ROIs in G; n = 4 mice/ genotype. Scale bar = 50 μm for low power images, 10 μm for insets in C and F. Scale bar in D = 1 μm in lower power TEM; 200 nm in inset TEM, as indicated. * *p* ≤ 0.05 by two-tailed Student’s t-test.

### Epithelial α3β1 integrin regulates organization of the alveolar BM

Having established a requirement for α3β1 integrin in BM component secretion in alveolar epithelial cells, we next investigated the organization of the alveolar BM. To assess the distance of the epithelium to the BM, we GFP-labeled the lung epithelium through generation of α3KO; mTmG and Shh Cre+; mTmG control mice, thereby marking α3-deficit or α3-replete Cre+ activated epithelium, respectively. We immunostained P28 lung sections from these mice for the BM marker nidogen-1 (magenta), a BM component that links collagen IV networks to laminin (42). As expected in control mTmG lungs, GFP+ epithelial cells co-localized with nidogen-1 along a thin alveolar septum (**Fig. 8A**). In contrast, α3KO; mTmG lungs contained areas where GFP-labeled epithelium failed to align with adjacent nidogen-1 (Yellow arrowheads in Fig. 8A), and in keeping with our TEM data, GFP+ epithelial cells were encased in the alveolar septum (asterisks). As this finding suggested significant disorganization of the BM, we examined the localization of nidogen-1 relative to laminin α5, the laminin isoform overabundant in the α3KO extracellular space (unlike laminin-332) (**Fig. 8B**). While laminin α5 co-localized with nidogen-1 along thin alveolar septa in control lungs, laminin α5 and nidogen-1 were disassociated from each other (arrowheads) in α3KO lungs. Since loss of α3β1 integrin disrupted both epithelial-nidogen and laminin-511/521-nidogen interactions, we next assessed laminin-collagen IV interactions in control and α3KO lungs at P28 by proximity ligation assay (PLA), which detects antibody-antigen interactions at ≤ 40 nm. Despite the relative overabundance of these isoforms in α3KO lungs (Fig. 5A, B), alveolar BMs from α3KO mice contained significantly fewer collagen IV α1-laminin α5 interactions at ≤ 40 nm than control lungs (**Fig. 8C**). These findings suggest alveolar epithelial cells require integrin α3β1 for BM component secretion and alveolar BM organization.

**Figure 8.**
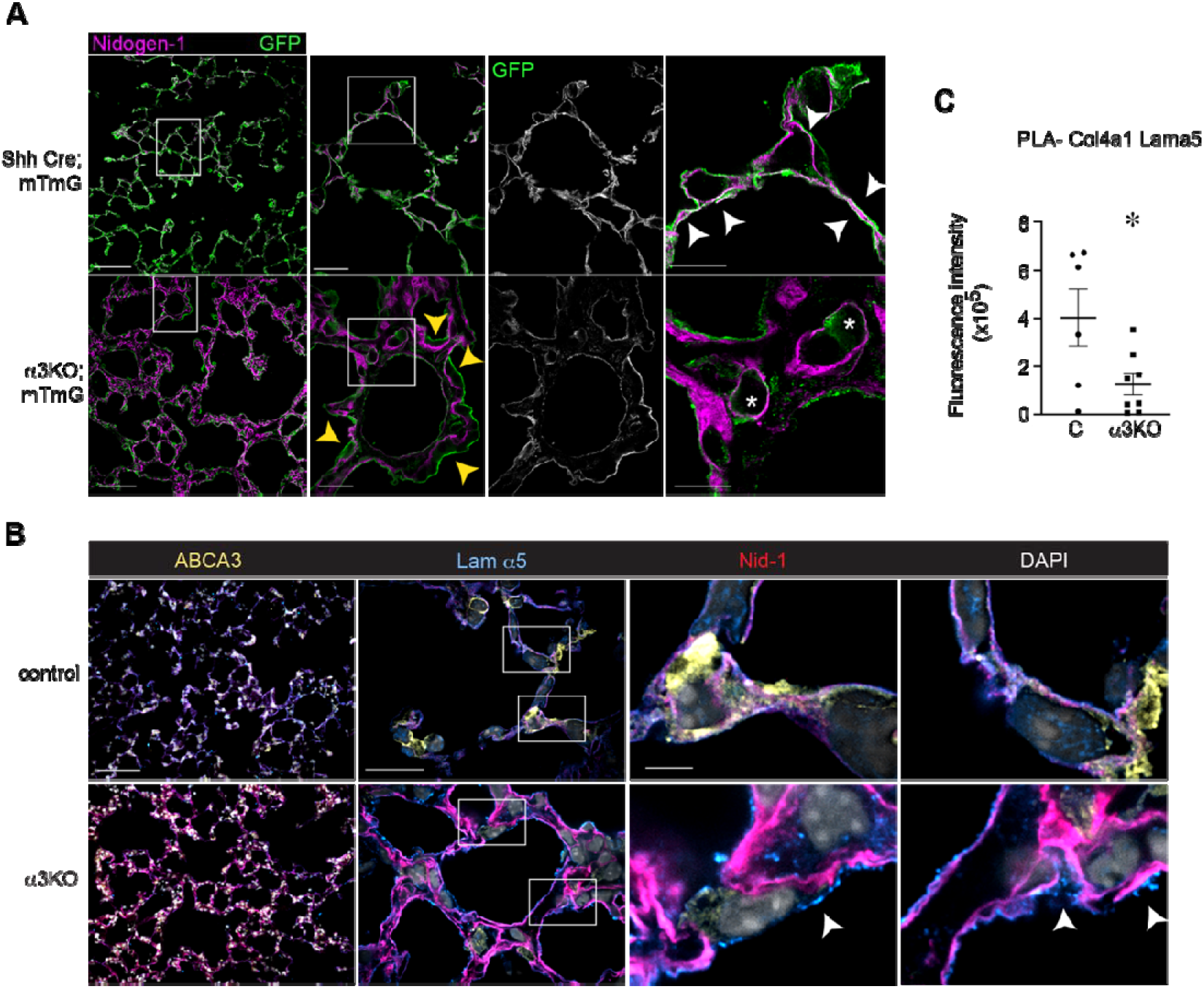
Integrin α3β1 regulates organization of the alveolar BM. (A) P28 α3KO; mTmG and Shh Cre+; mTmG immunostained for nidogen-1 (magenta); epithelium is labeled by GFP. White arrowheads mark areas of close epithelial-BM approximation in control lungs. Yellow arrowheads mark areas of epithelial-BM misalignment in α3KO; mTmG lungs. Asterisks = GFP+ epithelial cells within the alveolar septum in α3KO; mTmG lungs. (B) P28 α3KO and control lungs immunostained for ABCA3 (yellow), laminin α5 (blue), and nidogen-1 (red). DAPI nuclear stain in white. White arrowheads denote areas of laminin α5-nidogen-1 disassociation in α3KO lungs. (C) Proximity ligation assay of collagen IV α1 to laminin α5 demonstrates fewer points of proximity < 40 nm apart in α3KO lungs; n = 6 control and 8 α3KO lungs. Scale bar = 100 μm for low-power, 20 μm for high-power, 10 μm for inset in A; 100 μm for low-power, 25 μm for high-power, 10 μm for inset in B. * *p* ≤ 0.05 by Welch’s t-test.

## Discussion

The role of the epithelial LM-binding integrins in lung organogenesis is poorly understood. We demonstrate that integrin α3β1 is dispensable for airway branching but required for epithelial cell differentiation that is necessary for alveolar-capillary barrier formation during distal lung development. This unique role for α3β1 highlights the importance of precise epithelial-ECM interactions in lung development and challenges the paradigm that there is functional redundancy of LM-binding integrins in epithelial tissues. Additionally, our findings delineate requisite steps in the transition from branching morphogenesis into alveolarization, deepen our understanding of the influence of BM composition on epithelial cell behavior, and suggest that AT1 maturity is a requisite step in alveolar-capillary barrier function.

### Integrin α3β1-mediated epithelial differentiation directs alveolar-capillary barrier development

We report that alveolar epithelial differentiation is essential for alveolar BM formation through deposition of AT1-derived matrix proteins (16, 35, 43). Omission of AT1-specific BM components leaves the alveolar-capillary barrier structurally and functionally impaired, indicating that acquisition of these matrix components is required for maturation of alveolar-capillary barrier. Thus, our data establishes that epithelial differentiation is one of the initial steps in alveolar-capillary barrier development. Previously, we reported transcriptional evidence that a burst of epithelial cellular diversity at E18 precedes both endothelial and mesenchymal differentiation, which occurred from E18-P0 and P0-P5, respectively (34). Additionally, others have established a role for AT1-derived VEGF signaling that initiates capillary differentiation in the BM niche, adding evidence that AT1 maturation precedes endothelial differentiation within the developing alveolar-capillary barrier (1, 5, 10, 11). Thus, alveolar epithelial differentiation temporarily precedes and does not terminate until after the alveolar-capillary barrier is formed, bookending the critical epithelial-endothelial crosstalk essential for proper cellular organization.

Our findings suggest regulation of epithelial differentiation is a critical morphogenic program regulated by cell-matrix interactions in lung development. Previously, we reported that β1-containing integrins regulate alveolar epithelial differentiation in lung development and alveolar repair (23, 29) and now present evidence that integrin α3β1 performs a unique function. These findings suggest that integrins work in conjunction with other pathways, like YAP-Taz mechanotransduction pathways (44–50) and thyroid hormone signaling (51, 52), that potentiate alveolar epithelial cell differentiation. These studies address AT1 maturation in the context of development or alveolar regeneration; our work suggests an integrin-dependent role in development and additional investigation is needed to address the interaction of integrin α3β1 and these pathways.

### Precisely timed alveolar-capillary barrier formation allows for perinatal transition

Despite a well-established structural-functional relationship between alveolar-capillary barrier formation and gas exchange, the developmental programs that create a tight alveolar BM in normal mammalian lung development were previously unknown. Here, we defined the timing of normal epithelial-endothelial approximation, BM component isoform switching, and alveolar BM maturation during normal murine lung development. In the normal human and murine postnatal lung, the alveolar BM structure serves as a barrier, sealing airspace from interstitial fluid and preventing entry of external toxins and infectious agents into the lung. Our data indicate that the alveolar BM space is penetrable in the early perinatal period and the alveolar BM matures into a functional barrier postnatally, at P3, in the murine lung. We speculate that the delay in alveolar BM barrier function facilitates successful perinatal transition as secretion of fetal lung fluid into the developing airspaces supports lung growth, but this fluid must be rapidly reabsorbed back across the alveolar BM postnatally (53–55). It is possible that postnatal incorporation of collagen IV α3α4α5, whose numerous cysteine bonds thins the alveolar BM (43), allows the alveolar BM to fuse after the bulk of fetal lung fluid has been resorbed. Furthermore, this observation suggests that the mechanisms regulating fetal lung fluid clearance may be intimately tied to initiation of alveolar BM and/ or AT1 maturation. Consistent with this concept, alveolar epithelial differentiation and epithelial sodium channel receptors (ENaCs), which clear fetal lung fluid (56), are both regulated by glucocorticoid and thyroid hormone signaling (57–59).

### Limitations

This study focuses on the role of integrin α3β1 during distal lung development but does not address the mechanisms whereby 1) epithelial α3β1 regulates differentiation of endothelial cells in the alveolar-capillary barrier niche and 2) how integrin α3β1 expression by endothelial cells regulates formation of the alveolar-capillary barrier. While both these lines of investigation will likely result in substantial gains in mechanistic understanding of distal lung development, they are beyond the scope of this manuscript. Finally, the role of epithelial α3β1 in alveolar homeostasis and regeneration remains undefined and additional studies are needed to elucidate the timing and regulatory mechanisms in human lung development since term humans are born during alveolarization (60). Given the substantial impact of α3β1 on alveolar epithelial differentiation and alveolar BM formation during development, it is very likely that α3β1 regulates epithelial diversity and alveolar BM remodeling in the context of aging and injury, both in the murine and human lungs.

In conclusion, we report that during normal lung development, the ability to secrete specialized matrix components that tighten and refine the alveolar BM determines the threshold of AT1 cell functional maturity. Since α3β1 integrin regulates vesicle-based transport of matrix components, we show that α3β1 integrin is a key regulator of alveolar-capillary barrier formation and respiration.

## Methods

### Mice

We generated transgenic mice with pan-epithelial deletion of the α3 integrin subunit, termed α3KO mice, by crossing mice harboring the Shh-Cre allele (Jax#005622) (25, 26) with α3^flox/flox^ mice (27), generously gifted by Around Sonnenberg. Cre-littermate α3^f/f^ mice were used as controls (referred to as control mice). All mice were maintained on the C57BL/6 background, lungs were harvested at the indicated time points, and both male and female experimental mice were used in approximately equal proportions. All animal experiments were approved by the Institutional Animal Care and Use Committee at Vanderbilt University Medical Center.

### Histology and morphological analysis

Histological analysis was performed on hematoxylin and eosin-stained paraffin sections. For these experiments, mice were sacrificed at the indicated time points, the right ventricle was flushed with PBS, and inflation fixed at 25 cm with 10% formalin. Samples were paraffin processed, embedded, and sectioned at 8 μm thickness using standard techniques. Images were obtained on a Keyence BZ-X710 microscope. Immunofluorescence staining was performed on 8 μm frozen sections. Briefly, lungs were inflation fixed with a 2:1 PBS:O.C.T. (Tissue-Tek) mixture, embedded in O.C.T, and sectioned. Frozen section slides were then fixed with 4% paraformaldehyde, permeabilized with 0.1% Triton-X, incubated at 37 degrees in 5% donkey block for 2 hours at 37 degrees, incubated in primary antibody overnight at four degrees. Following secondary antibody incubation for 2 hours at room temperature, nuclei were DAPI stained as indicated, ProLong Gold mountant was applied, and imaged. Fluorescent images were obtained using a Nikon SoRa Spinning Disk Confocal with Super Resolution by Optical Pixel Reassignment (SoRa) super-resolution microscope equipped with a Hamamatsu ORCA-Fusion BT camera or on the Keyence BZ-X710 microscope. The following primary antibodies were used: anti-pro-SP-C (Abcam ab90716, 1:500); anti-ABCA3 (Abcam AB24751, 1:400); anti-T1α (Developmental Studies Hybridoma Bank 8.1.1, 1:400); anti-cytokeratin 8 (Origene BP5075, 1:100); anti-AGER (R&D AF1145, 1:200); anti-E-cadherin (Invitrogen 13-1900, 1:500); anti-integrin α3 (Invitrogen PA5-99407, 1:100); collagen IV α1 (Chondrex 7070, 1:100); collagen IV α3 (Chondrex 7076, 1:100); anti-laminin α5 (Kerafast EW1004, 1:200); anti-laminin β3 (Abcam ab97765, 1:100); anti-laminin γ2 (Invitrogen PA5-109901, 1:200); anti-nidogen-1 (Invitrogen MA1-06501, 1:200); anti-Rai14 (Proteintech 1507-1-AP, 1:200); anti-LL5B (Invitrogen PA5-113087, 1:200); and anti-GFP (Abcam ab13970, 1:200). For fluorescence activated cell sorting, we used anti-EpCAM-PE conjugated antibody (BD Biosciences 563477, 1:50) and anti-Ter119-APC conjugated antibody (BioLegend 116211, 1:200) for collection of EpCAM+ epithelial cells, along with PI viability dye (Sigma-Aldrich 537060, 1 μM). DAPI nuclear stain was used to mark nuclei (Cell Signaling 4083S, 1:10,000 in PBS). RNA *in situ* hybridization (RNAscope) for mouse *Sftpc*, *Hopx,* and *Mki67* (ACDBio) was performed per manufacturer’s instructions with inclusion of positive and negative control probes. Nonoverlapping images of RNAscope slides were obtained with a 40x objective (> 10 images/ mouse) and quantified. Proximity ligation assay was performed per manufacturer’s instructions. Fluorescence intensity measured on nonoverlapping 40x images, 15-20 images/ mouse.

### Ultrastructural examination

Lungs were harvested at the indicated timepoints, fixed with a 2% glutaraldehyde and 2% PFA solution in 0.1 M sodium cacodylate buffer, then sequentially post-fixed with 1% tannic acid, 1% OsO_4_, and en bloc stained with 1% uranyl acetate for 1 hour each. Samples were then dehydrated through a graded ethanol series and infiltrated with Epon-812 using propylene oxide as a transition solvent. The resin was polymerized at 60°C for 48 hours and thick sectioned at 1 μm as previously described (61). From the thick section, a 500 nm x 500 nm square region of interest was centered on an alveolar duct opening to determine the region for thin sectioning. Transmission electron microscopic images of the alveolar-capillary barrier, an area defined by an epithelial cell that directly overlies a capillary containing a single red blood cell, on a JEOL 2100 Plus Transmission Electron Microscope (200 keV LaB6 source) equipped with an AMT Nanosprint 15 MKII CMOS camera. Quantification of mean alveolar-capillary barrier width was calculated from perpendicular measurements obtained every 400 nm along the alveolar-capillary barrier.

### Functional testing of the alveolar-capillary barrier

Oxygen saturation was obtained by MouseOx probe on awake three-month-old α3KO and control mice, following recovery from light isoflurane inhalation anesthesia. Lung function testing was performed using the flexiVent apparatus (Scireq) on three-month-old α3KO and control mice following anesthetized with pentobarbital sodium (85 mg/kg)-induced anesthesia and 18-gauge tracheostomy. Mice were ventilated at standard parameters (150 breaths/ min and tidal volume = 10 ml/kg body weight), as previously reported (62). Arterial samples were obtained from three-month-old mice with in-dwelling arterial catheters breathing ambient room air (no supplemental oxygen administered). Analysis was performed on the i-STAT arterial blood gas analyzer per manufacturer’s instructions.

### Single-cell sample collection

Lung lobes were harvested from α3KO and control mice at P3 or between two and three months of age, minced, and incubated in dissociation media (RPMI-1640 with 0.7 mg/ml collagenase XI and 30 mg/ml type IV bovine pancreatic DNase) for 30 minutes at 37°C, as previously described (34). Disassociation into a single cell suspension was performed by passage through a wide bore pipet tip and filtration with a 40 μm filter, aliquoted, blocked with CD-32 Fc (BD cat# 553142) for 20 minutes on ice. Cells were washed with 2% fetal bovine serum staining buffer and incubated with anti-CD45 (BD cat #559864) and anti-Ter119 (BioLegend cat # 116211) conjugated flow cytometry antibodies. Samples generating the published WT transcriptomic data (34) were similarly processed at the indicated time points.

### scRNA-sequencing library preparation and next-generation sequencing

scRNA-seq libraries were generated in the Vanderbilt Technologies for Advanced Genomics Core (VANTAGE) using the Chromium Single Cell 5’ library kits (10X Genomics) and targeting 10,000-20,000 cells per sample, per the manufacturer’s instructions. Sequencing was performed on an Illumina NovaSeq 6000 instrument. For α3KO and control samples, CellRanger Count v7.1.0 (10X Genomics) was used to align reads to the mm10 reference genome. Published WT samples utilized CellRanger v3.1.

### P3 α3KO and control scRNA-sequencing analysis

P3 α3KO and control data were analyzed by the UNC Bioinformatics and Analytics Research Collaborative. Briefly, CellRanger filtered count matrices were loaded into R (v4.4.2) with Seurat (v5.3.1). Cells were demultiplexed with HTODemux. Cells were filtered to remove putative dead cells and background RNA (percent.mt ≤ 5, nCount_RNA ≥ 1200, nFeature_RNA ≥ 900). Data were normalized with SCTransform with each library as a batch variable and the percentage of mitochondrial RNAs as a regression variable. Libraries were then integrated using the Harmony method to mitigated batch-batch technical noise. Cells were annotated the ScType tool with the following marker genes to define alveolar subtypes. AT1: *Rtkn2, Aqp5, Ager, Col4a3*. AT2: *S*ftpa1*, Sftpc, Abca3, Lcn2*.

To isolate a clean alveolar subset, epithelial cells were re-normalized, re-clustered, and re-annotated to first extract AT2 and AT1 cells. Following isolation of clean AT2 and AT1 subsets, data were again re-normalized and clustered to produce the final alveolar population. The Seurat FindAllMarkers function with min.pct = 0.6 was used to identify top markers of unbiased alveolar clusters (resolution = 0.05). Manual inspection of the top marker *S100g* was used to identify the two clusters that comprised the Premature AT2 population. Plots were generated with Seurat, ggplot2, tidyHeatmap, and ComplexHeatmap.

### Adult scRNA-sequencing Analysis

Published WT and adult α3KO and control data were analyzed at Vanderbilt with a similar workflow as described above but with the following variations. Published WT data was analyzed as previously described (34). For adult α3KO and control data, analysis was performed by the Vanderbilt Creative Data Solutions Shared Resource. Analysis was performed in R (v4.1.1), using Seurat (v4.0.5) and SCTransform (v0.3.2.9008) with glmGamPoi (v1.6.0) (63–65). Cells were demultiplexed with HTODemux. Cells were filtered based on UMI threshold assessed by dropkick (66). Notably, data were not integrated since the average silhouette width (AWS) was close to zero, indicating that individual cells from the samples were thoroughly mixed and indistinguishable by technical origin. Gene counts for *Gm42418*, which is likely a rRNA, were removed (67). Identified by scDoubletFinder, 782 doublet cells were removed. Epithelial cells (*i.e., Epcam+* cell clusters) were sorted *in silico* for downstream analysis. Epithelial cells were annotated with manual inspection of the following marker genes: *Epcam*, *S*ftpa1, *Sftpc*, *Hopx*, *Aqp5*, *Col4a3*, *Ager*, *Foxj1*, *Dynlrb2*, *Mki67*, *Scgb1a1*, *Scgb3a2*, *Cdkn1a*, *Cldn4*, *Ascl1*, and *Scg5*.

Data integration was performed using a previously published dataset (68). Reads were downloaded from the Gene Expression Omnibus (GSE113049). CellRanger Count v3.1 with the mm10 genome was used for processing, followed by dimensionality reduction, clustering, and visualization as described above. Data were integrated with non-SoupX processed data using the ‘IntegrateData’ function in Seurat, following the workflow for data normalized with SCTransform. Samples from all experiments were combined, and clusters were annotated using marker genes described in the initial publication (68), and other canonical marker genes.

Cell label transfer was also utilized with a second published dataset (69). The annotated data matrix (h5ad file) was accessed as directed in the analysis code published as part of the methods (69). The raw counts from this annotated dataset were then processed using the Seurat workflow described above, while maintaining the published cell annotations. The Seurat “TransferData” function was then used to transfer the cell annotations onto the dataset generated here.

### Proteomics

Lungs harvested from P3 control and α3KO mice were homogenized, cell lysed, and fractionated into three fractions by solubility (intracellular, transmembrane, and extracellular matrix), with the extracellular fraction enriched for basement membrane proteins, per established methods (38). For our study, the intracellular and transmembrane were combined into a cellular fraction and the extracellular fraction was processed separately. Soluble samples were acetone precipitated overnight at -20°C. Proteins were pelleted, washed with cold acetone, resuspended in 5% SDS, 50 mM TEAB and agitated for 1.5 hours to ensure complete solubilization. Protein concentration was determined using a BCA assay. For each sample, 50 μg of protein were reduced with 5 mM DTT at 56 °C for 30 minutes, then alkylated with 15 mM IAA in the dark for 45 minutes. A final concentration of 2% phosphoric acid was then added to each sample, before loading onto S-trap microcolumns (Protifi KO2 micro). Protein digestion and cleaning were performed following the manufacturer’s protocol. Proteins were digested with trypsin (Promega #V15113) at a final concentration of 0.08 ug/ul. Peptides were eluted three times with 50 mM TEAB, then lyophilized. Samples were cleaned using C-18 spin columns (Thermo Scientific #PI89873). A fluorometric peptide assay was performed (Pierce #23275) to determine the peptide concentrations. Samples were dried down and stored at -80°C until further analysis. *LC/MS/MS Analysis:* All samples were reconstituted in 2% ACN in 0.1% formic acid and normalized to 0.1 µg/µl. Samples were analyzed via LC-MS/MS using a Vanquish Neo coupled to an Orbitrap Astral mass spectrometer (Thermo Scientific). A pooled sample was created by combining an aliquot of each sample, which was analyzed intermittently throughout the sequence to assess technical reproducibility. Samples (2 µl) were injected onto an Aurora Ultimate TS column (75 μm id × 25 cm, 1.7 μm particle size; IonOpticks) and separated over a 30 min method. The gradient consisted of 5-45% mobile phase B at a 300 nl/min flow rate, where mobile phase A was 0.1% formic acid in water and mobile phase B was 0.1% formic acid in 80% acetonitrile. MS1 full scans (m/z 350-980) were acquired with a resolution of 240,000 in the Orbitrap analyzer with a maximum injection time of 5 ms and an AGC target of 500%. MS2 scans were acquired using the Astral detector operated in Data-Independent Acquisition (DIA) mode, covering a scan range of m/z 150-2000 with a cycle time of 0.6 s. Isolation windows were set to 3 m/z with a higher collision dissociation (HCD) energy of 25%, an AGC target of 500%, and a maximum injection time of 3 ms.

### Data Analysis

Raw data files were processed using Spectronaut (v20; Biognosys) and searched against the Uniprot reviewed mouse database (UP000000589, containing 17,230 entries, downloaded January 2025) and the Hao Lab mouse tissue common contaminants database (277 entries). We used the following parameters: enzyme specificity set to trypsin, up to two missed cleavages allowed, cysteine carbamidomethylation set as a fixed modification, methionine oxidation and N-terminal acetylation set as variable modifications. A false discovery rate (FDR) of 1% was used to filter all data. Imputation was disabled, and single hit proteins were retained. The mass spectrometry proteomics data have been deposited to the ProteomeXchange Consortium via the PRIDE [1] partner repository with the dataset identifier PXDXXXXXX.

### Statistics

Comparison between two groups was performed by two-tailed t-test as indicated using Prism GraphPad statistical software. Individual *p-*values, q-values, and sample size for each group are included in the figure legends. For graphical data, data represent mean ± SEM. ScRNA-sequencing analysis was completed as above.

## Data availability

A complete collection of all package versions, and code for all steps of the WT transcriptional analysis is available at https://github.com/SucreLab. All sequencing data have been deposited to the NCBI GEO database with accession number GSEXXXXXX. Proteomics data has been deposited in PRIDE, number PXDXXXXXX. Other data available upon request.

## Author contributions

Conceptualization: TSB, RZ, EJP; Methodology: JMSS, JPC, NB, RH, JJR, AS, EJP; Validation: JJG, JMSS, EJP; Formal analysis: MJN, AKY, NRM, ALM, AJH, SPS, TSB, RZ, EJP; Investigation: JJG, ALS, DCN, EJP; Data curation: ADH, ALA, AKY, NRM, ALM, AJH, ALS, NB, SK, JCL, CSJ, DCN, RH, EJP; Writing-original draft: MJN, RZ, EJP; Writing-review and editing: MJN, JMSS, TSB, RZ, EJP; Visualization: MJN, ALM, AJH, JPC, EJP; Supervision: EJP; Project administration: AJH, EJP; Funding acquisition: EJP

## Acknowledgements

The authors would like to thank David H. Wasserman, Whitney Longmate, Michael DiPersio, Nathan C. Winn, Ambra Pozzi, Jonathan Kropski, Angela Jones, Evan Krystofiak, and Kristie Rose for content expertise and David Nichols for technical assistance. This work was supported by NIH grants R03 HL 154287 (EJP), R01 HL163195 (EJP), R01 HL176912 (JJG), R01 HL164327 (JJR), and R01 HL175555 (TSB), as well as VA Merit Review 2 l01 BX002378 (TSB). JJG is supported by the Parker B. Francis Family Foundation. RL is funded by the Wellcome Trust (Ref: 226804/Z/22/Z, Ref: 227417/Z/23/Z, Ref: 301803/Z/23/Z); Kidney Research UK and The Stoneygate Trust (Alport Research Hub); the National Institute for Health Research (NIHR-Manchester BRC); FourPoints Innovation, a portfolio company of certain funds managed by Deerfield Management Company; Visterra Therapeutics and Ascidian Therapeutics. She has consultancy agreements with Calliditas Therapeutics and Travere Therapeutics. Transmission electron and confocal microscopy was performed in part utilizing the Vanderbilt Cell Imaging Shared Resource (supported by NIH grants CA68485, DK20593, DK58404, DK59637, EY08126, 1S10OD034315-01, 1R24OD037694-01) and the UNC Hooker Imaging Core Facility (supported by P30 CA016086 Cancer Center Core Support Grant to the UNC Lineberger Comprehensive Cancer Center). ScRNA-sequencing experiments were performed using the Vanderbilt Technologies for Advanced Genomics (VANTAGE) Core facility. The Vanderbilt Creative Data Solutions Shared Resource (RRID:SCR_022366) performed the analysis of the adult α3 scRNA-sequencing experiments. P3 scRNA-sequencing data was analyzed through the Bioinformatics and Analytics Research Collaborative (BARC) at UNC. Flow cytometry experiments were performed in the VUMC Flow Cytometry Shared Resource (supported by the Vanderbilt Ingram Cancer Center P30 CA68485 and the Vanderbilt Digestive Disease Research Center DK058404). *This research is based in part upon work conducted using the UNC Michael Hooker Metabolomics and Proteomics Core Facility, which is supported by the NCI Cancer Center Support Grant to the UNC Lineberger Comprehensive Cancer Center, P30 CA016086, and the UNC Nutrition Obesity Research Center, P30 DK056350.* Operations in the Pulmonary Inhalation Research Core is supported in part by the Cystic Fibrosis Foundation Research and Development Program (ESTHER24R0) and the NIDDK Cystic Fibrosis Research and Translation Core Center (P30 DK065988-21).

## References

1. Vila Ellis L, Cain MP, Hutchison V, Flodby P, Crandall ED, Borok Z, et al. Epithelial Vegfa Specifies a Distinct Endothelial Population in the Mouse Lung. Dev Cell. 2020;52(5):617–30 e6.

2. Schittny JC. Development of the lung. Cell Tissue Res. 2017;367(3):427–44.

3. Frank DB, Penkala IJ, Zepp JA, Sivakumar A, Linares-Saldana R, Zacharias WJ, et al. Early lineage specification defines alveolar epithelial ontogeny in the murine lung. Proc Natl Acad Sci U S A. 2019;116(10):4362–71.

4. Rao S, Liu M, Iosef C, Knutsen C, and Alvira CM. Endothelial-specific loss of IKKbeta disrupts pulmonary endothelial angiogenesis and impairs postnatal lung growth. Am J Physiol Lung Cell Mol Physiol. 2023;325(3):L299–L313.

5. Zanini F, Che X, Knutsen C, Liu M, Suresh NE, Domingo-Gonzalez R, et al. Developmental diversity and unique sensitivity to injury of lung endothelial subtypes during postnatal growth. iScience. 2023;26(3):106097.

6. Gillich A, Zhang F, Farmer CG, Travaglini KJ, Tan SY, Gu M, et al. Capillary cell-type specialization in the alveolus. Nature. 2020;586(7831):785–9.

7. Negretti NM, Son Y, Crooke P, Plosa EJ, Benjamin JT, Jetter CS, et al. Epithelial outgrowth through mesenchymal rings drives lung alveologenesis. JCI Insight. 2025;10(4).

8. Frank DB, Peng T, Zepp JA, Snitow M, Vincent TL, Penkala IJ, et al. Emergence of a Wave of Wnt Signaling that Regulates Lung Alveologenesis by Controlling Epithelial Self-Renewal and Differentiation. Cell Rep. 2016;17(9):2312–25.

9. Whitsett JA, Kalin TV, Xu Y, and Kalinichenko VV. Building and Regenerating the Lung Cell by Cell. Physiol Rev. 2019;99(1):513–54.

10. Niethamer TK, Stabler CT, Leach JP, Zepp JA, Morley MP, Babu A, et al. Defining the role of pulmonary endothelial cell heterogeneity in the response to acute lung injury. Elife. 2020;9.

11. Yang J, Hernandez BJ, Martinez Alanis D, Narvaez del Pilar O, Vila-Ellis L, Akiyama H, et al. The development and plasticity of alveolar type 1 cells. Development. 2016;143(1):54–65.

12. Timpl R, and Brown JC. Supramolecular assembly of basement membranes. Bioessays. 1996;18(2):123–32.

13. Yurchenco PD. Basement membranes: cell scaffoldings and signaling platforms. Cold Spring Harb Perspect Biol. 2011;3(2).

14. Keeley DP, and Sherwood DR. Tissue linkage through adjoining basement membranes: The long and the short term of it. Matrix Biol. 2019;75–76:58–71.

15. Pozzi A, Yurchenco PD, and Iozzo RV. The nature and biology of basement membranes. Matrix Biol. 2017;57–58:1–11.

16. Abrahamson DR, St John PL, Stroganova L, Zelenchuk A, and Steenhard BM. Laminin and type IV collagen isoform substitutions occur in temporally and spatially distinct patterns in developing kidney glomerular basement membranes. J Histochem Cytochem. 2013;61(10):706–18.

17. Coraux C, Delplanque A, Hinnrasky J, Peault B, Puchelle E, and Gaillard D. Distribution of integrins during human fetal lung development. J Histochem Cytochem. 1998;46(7):803–10.

18. Hynes RO. Integrins: bidirectional, allosteric signaling machines. Cell. 2002;110(6):673–87.

19. Has C, Sparta G, Kiritsi D, Weibel L, Moeller A, Vega-Warner V, et al. Integrin alpha3 mutations with kidney, lung, and skin disease. N Engl J Med. 2012;366(16):1508–14.

20. Nicolaou N, Margadant C, Kevelam SH, Lilien MR, Oosterveld MJ, Kreft M, et al. Gain of glycosylation in integrin alpha3 causes lung disease and nephrotic syndrome. J Clin Invest. 2012;122(12):4375–87.

21. Shukrun R, Vivante A, Pleniceanu O, Vax E, Anikster Y, Dekel B, et al. A human integrin-alpha3 mutation confers major renal developmental defects. PLoS One. 2014;9(3):e90879.

22. Kreidberg JA, Donovan MJ, Goldstein SL, Rennke H, Shepherd K, Jones RC, et al. Alpha 3 beta 1 integrin has a crucial role in kidney and lung organogenesis. Development. 1996;122(11):3537–47.

23. Plosa EJ, Young LR, Gulleman PM, Polosukhin VV, Zaynagetdinov R, Benjamin JT, et al. Epithelial beta1 integrin is required for lung branching morphogenesis and alveolarization. Development. 2014;141(24):4751–62.

24. Kim KK, Wei Y, Szekeres C, Kugler MC, Wolters PJ, Hill ML, et al. Epithelial cell alpha3beta1 integrin links beta-catenin and Smad signaling to promote myofibroblast formation and pulmonary fibrosis. J Clin Invest. 2009;119(1):213– 24.

25. Harfe BD, Scherz PJ, Nissim S, Tian H, McMahon AP, and Tabin CJ. Evidence for an expansion-based temporal Shh gradient in specifying vertebrate digit identities. Cell. 2004;118(4):517–28.

26. Harris KS, Zhang Z, McManus MT, Harfe BD, and Sun X. Dicer function is essential for lung epithelium morphogenesis. Proc Natl Acad Sci U S A. 2006;103(7):2208–13.

27. Sachs N, Kreft M, van den Bergh Weerman MA, Beynon AJ, Peters TA, Weening JJ, et al. Kidney failure in mice lacking the tetraspanin CD151. J Cell Biol. 2006;175(1):33–9.

28. Chen J, and Krasnow MA. Integrin Beta 1 suppresses multilayering of a simple epithelium. PLoS One. 2012;7(12):e52886.

29. Sucre JM, Bock F, Negretti NM, Benjamin JT, Gulleman PM, Dong X, et al. Alveolar repair following LPS-induced injury requires cell-ECM interactions. JCI Insight. 2023;8(14).

30. Dupret JM, L’Horset F, Perret C, Bernaudin JF, and Thomasset M. Calbindin-D9K gene expression in the lung of the rat. Absence of regulation by 1,25-dihydroxyvitamin D3 and estrogen. Endocrinology. 1992;131(6):2643–8.

31. Brownfield DG, de Arce AD, Ghelfi E, Gillich A, Desai TJ, and Krasnow MA. Alveolar cell fate selection and lifelong maintenance of AT2 cells by FGF signaling. Nat Commun. 2022;13(1):7137.

32. Treutlein B, Brownfield DG, Wu AR, Neff NF, Mantalas GL, Espinoza FH, et al. Reconstructing lineage hierarchies of the distal lung epithelium using single-cell RNA-seq. Nature. 2014;509(7500):371–5.

33. Zepp JA, Morley MP, Loebel C, Kremp MM, Chaudhry FN, Basil MC, et al. Genomic, epigenomic, and biophysical cues controlling the emergence of the lung alveolus. Science. 2021;371(6534).

34. Negretti NM, Plosa EJ, Benjamin JT, Schuler BA, Habermann AC, Jetter CS, et al. A single-cell atlas of mouse lung development. Development. 2021;148(24).

35. Abrahamson DR, Hudson BG, Stroganova L, Borza DB, and St John PL. Cellular origins of type IV collagen networks in developing glomeruli. J Am Soc Nephrol. 2009;20(7):1471–9.

36. Miner JH, Patton BL, Lentz SI, Gilbert DJ, Snider WD, Jenkins NA, et al. The laminin alpha chains: expression, developmental transitions, and chromosomal locations of alpha1-5, identification of heterotrimeric laminins 8-11, and cloning of a novel alpha3 isoform. J Cell Biol. 1997;137(3):685–701.

37. Miner JH, and Sanes JR. Collagen IV alpha 3, alpha 4, and alpha 5 chains in rodent basal laminae: sequence, distribution, association with laminins, and developmental switches. J Cell Biol. 1994;127(3):879–91.

38. Morais M, Tian P, O’Cualain R, Lawless C, and Lennon R. Protocol to characterize basement membranes during kidney development using mass spectrometry-based label-free quantitative proteomics. STAR Protoc. 2023;4(4):102741.

39. Naba A, Clauser KR, Hoersch S, Liu H, Carr SA, and Hynes RO. The matrisome: in silico definition and in vivo characterization by proteomics of normal and tumor extracellular matrices. Mol Cell Proteomics. 2012;11(4):M111 014647.

40. Chioran A, Duncan S, Catalano A, Brown TJ, and Ringuette MJ. Collagen IV trafficking: The inside-out and beyond story. Dev Biol. 2017;431(2):124–33.

41. Hotta A, Kawakatsu T, Nakatani T, Sato T, Matsui C, Sukezane T, et al. Laminin-based cell adhesion anchors microtubule plus ends to the epithelial cell basal cortex through LL5alpha/beta. J Cell Biol. 2010;189(5):901–17.

42. Aumailley M, Wiedemann H, Mann K, and Timpl R. Binding of nidogen and the laminin-nidogen complex to basement membrane collagen type IV. Eur J Biochem. 1989;184(1):241–8.

43. Pokidysheva EN, Redhair N, Ailsworth O, Page-McCaw P, Rollins-Smith L, Jamwal VS, et al. Collagen IV of basement membranes: II. Emergence of collagen IV(alpha345) enabled the assembly of a compact GBM as an ultrafilter in mammalian kidneys. J Biol Chem. 2023;299(12):105459.

44. DiGiovanni GT, Han W, Sherrill TP, Taylor CJ, Nichols DS, Geis NM, et al. Epithelial Yap/Taz are required for functional alveolar regeneration following acute lung injury. JCI Insight. 2023;8(19).

45. Gaona IP, McCall AS, Geis NM, Colvard AC, DiGiovanni GT, Sherrill TP, et al. Sustained Yap/Taz activation promotes aberrant alveolar epithelial cell differentiation and drives persistent fibrotic remodeling. JCI Insight. 2026.

46. Gokey JJ, Snowball J, Sridharan A, Sudha P, Kitzmiller JA, Xu Y, et al. YAP regulates alveolar epithelial cell differentiation and AGER via NFIB/KLF5/NKX2-1. iScience. 2021;24(9):102967.

47. LaCanna R, Liccardo D, Zhang P, Tragesser L, Wang Y, Cao T, et al. Yap/Taz regulate alveolar regeneration and resolution of lung inflammation. J Clin Invest. 2019;129(5):2107–22.

48. Volckaert T, Yuan T, Yuan J, Boateng E, Hopkins S, Zhang JS, et al. Hippo signaling promotes lung epithelial lineage commitment by curbing Fgf10 and beta-catenin signaling. Development. 2019;146(2).

49. Burgess CL, Huang J, Bawa PS, Alysandratos KD, Minakin K, Ayers LJ, et al. Generation of human alveolar epithelial type I cells from pluripotent stem cells. Cell Stem Cell. 2024;31(5):657–75 e8.

50. Nantie LB, Young RE, Paltzer WG, Zhang Y, Johnson RL, Verheyden JM, et al. Lats1/2 inactivation reveals Hippo function in alveolar type I cell differentiation during lung transition to air breathing. Development. 2018;145(21).

51. Pan X, Wang L, Yang J, Li Y, Xu M, Liang C, et al. TRbeta activation confers AT2-to-AT1 cell differentiation and anti-fibrosis during lung repair via KLF2 and CEBPA. Nat Commun. 2024;15(1):8672.

52. Wang L, Li Z, Wan R, Pan X, Li B, Zhao H, et al. Single-Cell RNA Sequencing Provides New Insights into Therapeutic Roles of Thyroid Hormone in Idiopathic Pulmonary Fibrosis. Am J Respir Cell Mol Biol. 2023;69(4):456–69.

53. Adamson TM, Boyd RD, Platt HS, and Strang LB. Composition of alveolar liquid in the foetal lamb. J Physiol. 1969;204(1):159–68.

54. Aherne W, and Dawkins MJ. The Removal of Fluid from the Pulmonary Airways after Birth in the Rabbit, and the Effect on This of Prematurity and Pre-Natal Hypoxia. Biol Neonat. 1964;7:214–29.

55. Olver RE, and Strang LB. Ion fluxes across the pulmonary epithelium and the secretion of lung liquid in the foetal lamb. J Physiol. 1974;241(2):327–57.

56. Smith DE, Otulakowski G, Yeger H, Post M, Cutz E, and O’Brodovich HM. Epithelial Na(+) channel (ENaC) expression in the developing normal and abnormal human perinatal lung. Am J Respir Crit Care Med. 2000;161(4 Pt 1):1322–31.

57. Gonzales LW, Guttentag SH, Wade KC, Postle AD, and Ballard PL. Differentiation of human pulmonary type II cells in vitro by glucocorticoid plus cAMP. Am J Physiol Lung Cell Mol Physiol. 2002;283(5):L940–51.

58. Otulakowski G, Rafii B, Harris M, and O’Brodovich H. Oxygen and glucocorticoids modulate alphaENaC mRNA translation in fetal distal lung epithelium. Am J Respir Cell Mol Biol. 2006;34(2):204–12.

59. Richard K, Ramminger SJ, Forsyth L, Burchell A, and Wilson SM. Thyroid hormone potentiates glucocorticoid-evoked airway Na+ transport without affecting alpha-ENaC transcription. FEBS Lett. 2004;576(3):339–42.

60. Pan H, Deutsch GH, Wert SE, Ontology S, and Consortium NMAoLDP. Comprehensive anatomic ontologies for lung development: A comparison of alveolar formation and maturation within mouse and human lung. J Biomed Semantics. 2019;10(1):18.

61. Plosa EJ, Benjamin JT, Sucre JM, Gulleman PM, Gleaves LA, Han W, et al. beta1 Integrin regulates adult lung alveolar epithelial cell inflammation. JCI Insight. 2020;5(2).

62. Gutor SS, Salinas RI, Nichols DS, Bazzano JMR, Han W, Gokey JJ, et al. Repetitive sulfur dioxide exposure in mice models post-deployment respiratory syndrome. Am J Physiol Lung Cell Mol Physiol. 2024;326(5):L539–L50.

63. Hao Y, Hao S, Andersen-Nissen E, Mauck WM, 3rd, Zheng S, Butler A, et al. Integrated analysis of multimodal single-cell data. Cell. 2021;184(13):3573–87 e29.

64. Hafemeister C, and Satija R. Normalization and variance stabilization of single-cell RNA-seq data using regularized negative binomial regression. Genome Biol. 2019;20(1):296.

65. Ahlmann-Eltze C, and Huber W. glmGamPoi: fitting Gamma-Poisson generalized linear models on single cell count data. Bioinformatics. 2021;36(24):5701–2.

66. Heiser CN, Wang VM, Chen B, Hughey JJ, and Lau KS. Automated quality control and cell identification of droplet-based single-cell data using dropkick. Genome Res. 2021;31(10):1742–52.

67. Kimmel JC, Hwang AB, Scaramozza A, Marshall WF, and Brack AS. Aging induces aberrant state transition kinetics in murine muscle stem cells. Development. 2020;147(9).

68. Riemondy KA, Jansing NL, Jiang P, Redente EF, Gillen AE, Fu R, et al. Single cell RNA sequencing identifies TGFbeta as a key regenerative cue following LPS-induced lung injury. JCI Insight. 2019;5.

69. Strunz M, Simon LM, Ansari M, Kathiriya JJ, Angelidis I, Mayr CH, et al. Alveolar regeneration through a Krt8+ transitional stem cell state that persists in human lung fibrosis. Nat Commun. 2020;11(1):3559.

