## Supplemental figures for "Integrin α3β1 is required for alveolar-capillary barrier formation through regulation of epithelial differentiation"

Supplemental Figure 1

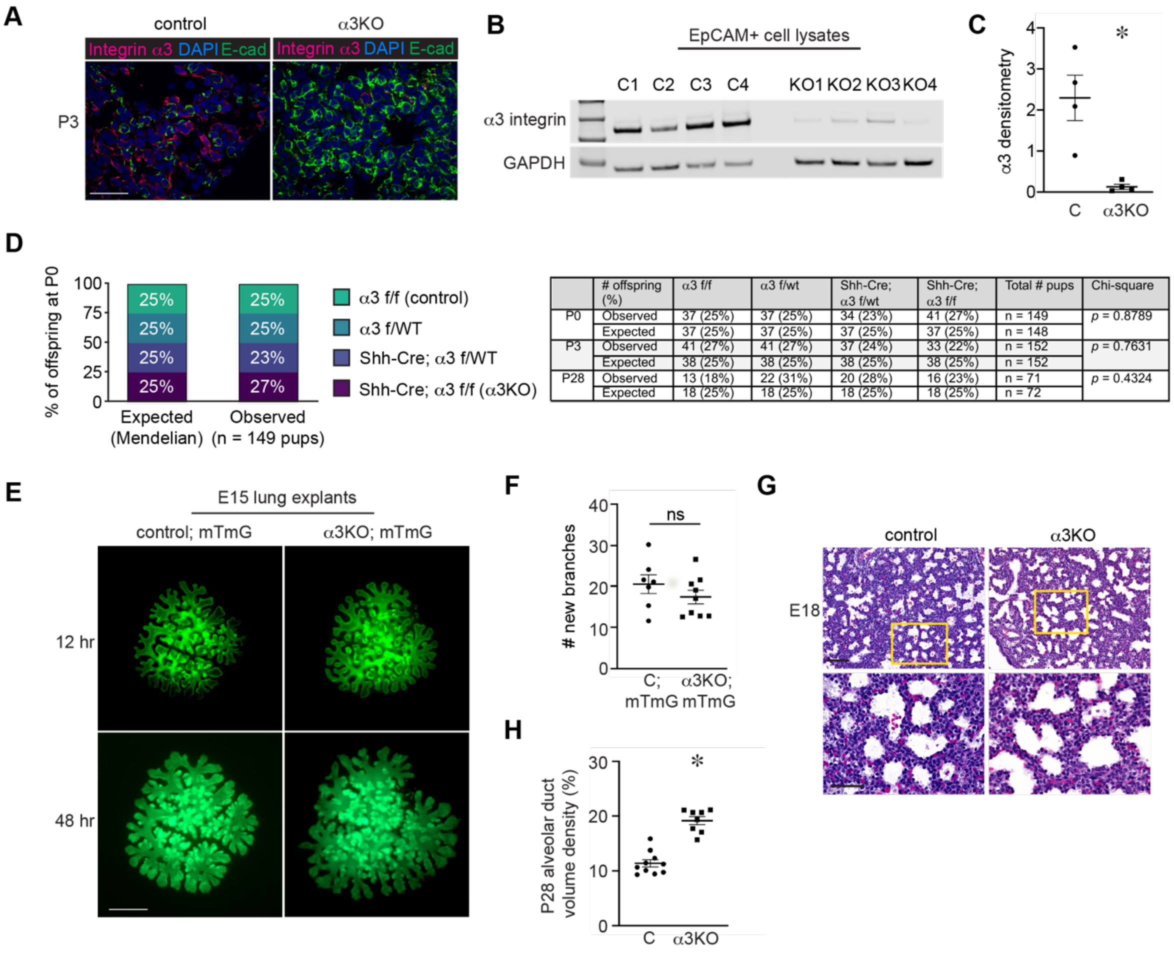


| | # offspring (%) | α3 f/f | α3 f/wt | Shh-Cre; α3 f/wt | Shh-Cre; α3 f/f | Total # pups | Chi-square |
|---|---|---|---|---|---|---|---|
| P0 | Observed | 37 (25%) | 37 (25%) | 34 (23%) | 41 (27%) | n = 149 | *p* = 0.8789 |
| | Expected | 37 (25%) | 37 (25%) | 37 (25%) | 37 (25%) | n = 148 | |
| P3 | Observed | 41 (27%) | 41 (27%) | 37 (24%) | 33 (22%) | n = 152 | *p* = 0.7631 |
| | Expected | 38 (25%) | 38 (25%) | 38 (25%) | 38 (25%) | n = 152 | |
| P28 | Observed | 13 (18%) | 22 (31%) | 20 (28%) | 16 (23%) | n = 71 | *p* = 0.4324 |
| | Expected | 18 (25%) | 18 (25%) | 18 (25%) | 18 (25%) | n = 72 | |

**Figure S1.** (A) Immunostained lung sections for the integrin α3 subunit (magenta) and E-cadherin (green) demonstrate α3 epithelial-specific deletion in P3 α3KO lungs. (B) Immunoblot shows near absence of the α3 subunit in P28 α3KO EpCAM+ primary epithelial cell lysates, n = 4 mice/ genotype. (C) Immunoblot densitometry standardized to GAPDH for B. (D) Expected and observed genotypes from α3KO breeding strategy. Genotype frequency by Chi-square analysis demonstrates α3KO mice are born in normal Mendelian ratio. (E) Fluorescence imaging of E15 lung explants generated from α3KO and control mice crossed to the mTmG reporter demonstrates no difference in the number of new peripheral branches from 12 to 48 hours, quantified in (F), (17.3±1.6 branches in α3KO explants vs. 20.5±2.3 branches in control explants; n = 7 control explants and 9 α3KO explants). (G) Images of hematoxylin and eosin-stained lung sections from E18 α3KO and control lungs show similar gross morphology at the end of sacculation. Scale bar = 50 μm for A, 500 μm for E, 100 μm for G. (H) Increased alveolar duct volume density in P28 α3KO lungs compared to control lungs; 11.4±0.7 μm in control lungs vs. 19.16±0.7 μm in α3KO lungs; n = 10 control and 8 α3KO lungs. * $p \leq 0.05$ for C by Welch's t test.

Supplemental Figure 2

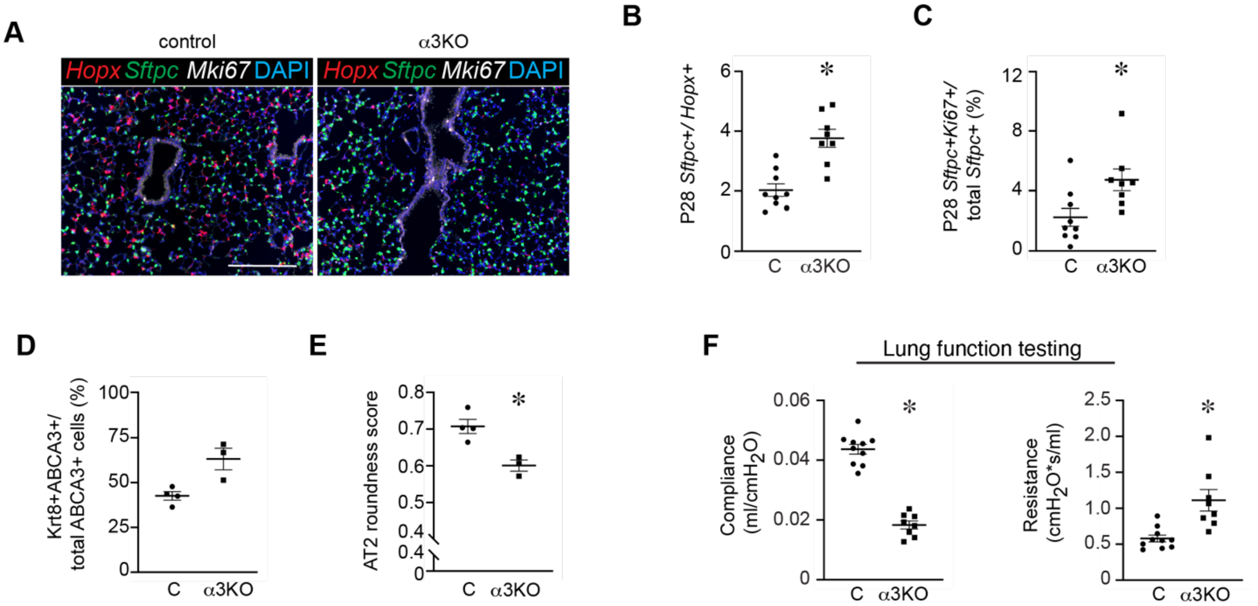


**Figure S2.** (A) RNA in situ hybridization (RNAscope) for *Sftpc* (green)*, Hopx* (red)*,* and the proliferation marker *Mki67* (white) with DAPI nuclear stain (blue) shows an overabundance of AT2 cells and a paucity of AT1 cells in P28 α3KO lungs, quantified by an increased AT2/ AT1 ratio (B). (C) Increased AT2 proliferation in P28 α3KO lungs, 3.8±0.3% for α3KO lungs vs. 2.0±0.2% for control lungs. For B and C, n = 9 control and 8 α3KO lungs; ten 40x sections/ mouse. (D) Nonsignificant increase in dual ABCA3+/ Krt8+ epithelial cells in α3KO lungs compared to control lungs, 42.6±2.4% in control lungs vs. 63.2±6.1% in α3KO lungs, n = 4 control and 3 α3KO lungs, ten 20x sections/ mouse, $p = 0.23$ by Welch's t test. (E) Decreased roundness score for ABCA3+ epithelial cells in α3KO lung compared to control lungs, 0.71±0.02 roundness score in control lungs vs. 0.60±0.02, n = 4 control and 3 α3KO lungs, 54-90 cells/ mouse analyzed. (F) FlexiVent lung function testing demonstrated decreased compliance (0.183±0.001 ml/cm$H_2O$ for α3KO vs. 0.044±0.002 ml/cm$H_2O$ for control mice) and increased resistance (1.11±0.15 ml/cm$H_2O$ for α3KO vs. 0.58±0.05 ml/cm$H_2O$ for control mice) (n = 10 control mice, 8 α3KO mice). $* \, p < 0.05$ by Welch's t test for B, C, E, and F.

Supplemental Figure 3

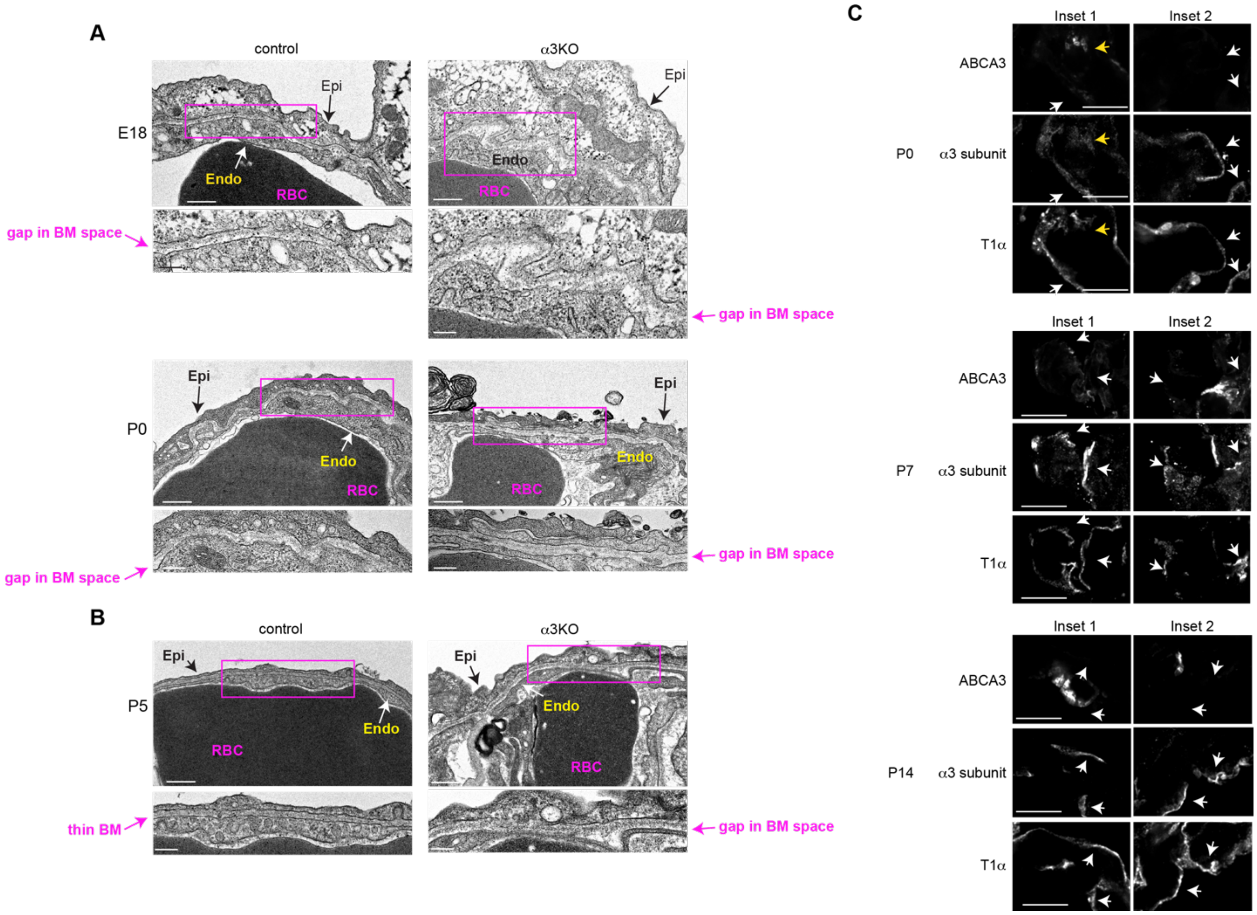


**Figure S3**. (A) Transmission electron micrographs (TEM) of control and α3KO lungs showing incomplete alveolar BM formation at E18 and P0. (B) By P5, control alveolar BMs have matured into a thin matrix structure, whereas α3KO alveolar BM is discontinuous with gaps in the matrix and a widened BM space between endothelial and epithelial cells. Scale bar = 500 nm for low-power TEM (top panels), 200 nm for TEM insets (bottom panels) in A and B. Epi = epithelial cell, Endo = capillary endothelial cell, RBC = red blood cell. (C) Single channels for images of P0, P7, and P14 WT lung immunostained for ABCA3, the α3 integrin subunit, and T1α, as indicated. Scale bar = 5 µm for high-power insets in C.

Supplemental Figure 4

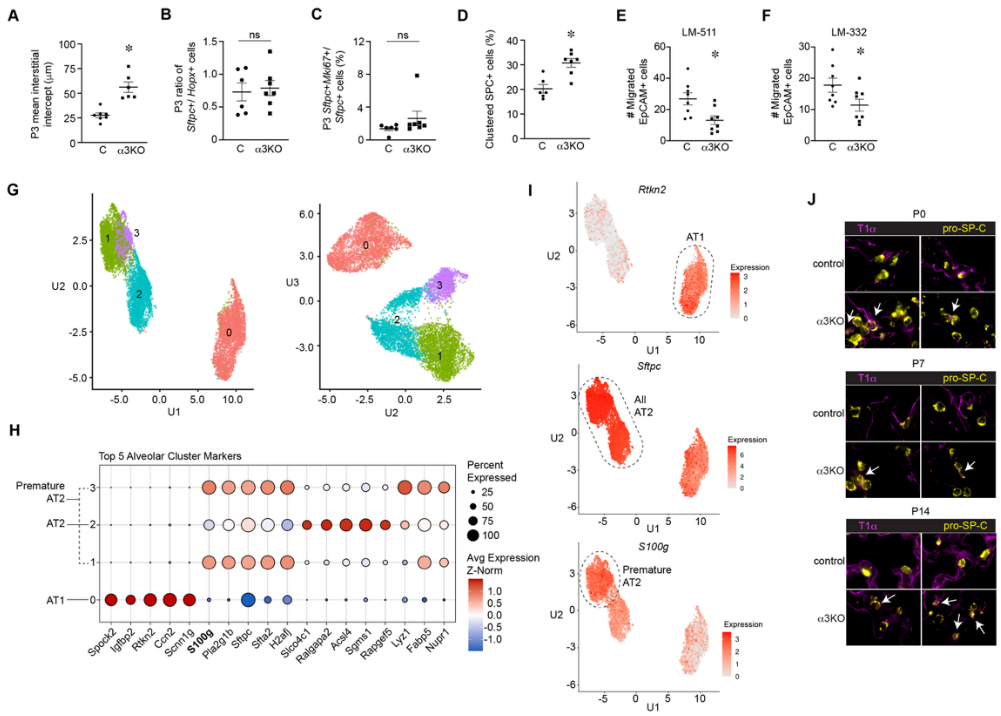


**Supp Fig 4.** (A) Increased mean interstitial intercept in P3 α3KO lungs compared to control P3 lungs, 56.1±5.3 μm for α3KO lungs vs. 27.6±2.3 μm for control lungs (n = 7 control, 6 α3KO mice). (B) No difference in the ratio of *Sftpc+/ Hopx+* cells or the percentage of proliferating AT2 cells between the genotypes at P3. (C) No difference in AT2 proliferation in P3 α3KO and control lungs; 3.8±0.3% for α3KO lungs vs. 2.0±0.2% for control lungs, $p = 0.21$. For B and C, cells identified by RNAscope probes for *Sftpc+, Hopx+,* and the proliferation marker *Mki67*. (n = 6 control, 7 α3KO mice). (D) Increased percentage of *Sftpc+* epithelial cells cluster in pairs or triplets at P3 in α3KO lungs compared to control P3 lungs, 20.3±1.7% in P3 control lungs vs. 30.8±1.8% in P3 α3KO lungs, n = 6 control, 7 α3KO mice). (E-F) Migration assay for EpCAM+ epithelial cells collected by FACS across Transwell filters in 6 hours on laminin-511 (E) and laminin-332 (F). For laminin-511, 26.8±3.9 migrated control cells vs. 13.1±2.8 migrated α3KO cells. For laminin-332, 17.8±2.3 migrated control cells vs. 11.4±1.9 migrated α3KO cells. In both (E) and (F), assays performed in triplicate with cells collected from n= 8 mice/ genotype. (G-H) UMAP showing 3 distinct alveolar epithelial cell clusters at P3, annotated by unbiased cluster-defining differential gene signatures in H. (I) UMAP representation of hallmark gene expression for AT1 (*Rtkn2)*, AT2 (*Sftpc*), and premature AT2 cells (*S100g*). For G-I, n = 5 control mice, 8 α3KO mice. (J) Control and α3KO lungs immunostained for T1α (magenta) and pro-SP-C (yellow). White arrows denote dual T1α+/ pro-SP-C+ cells. ∗ $p < 0.05$ by Welch's t test for A, D, E, and F.

Supplemental Figure 5

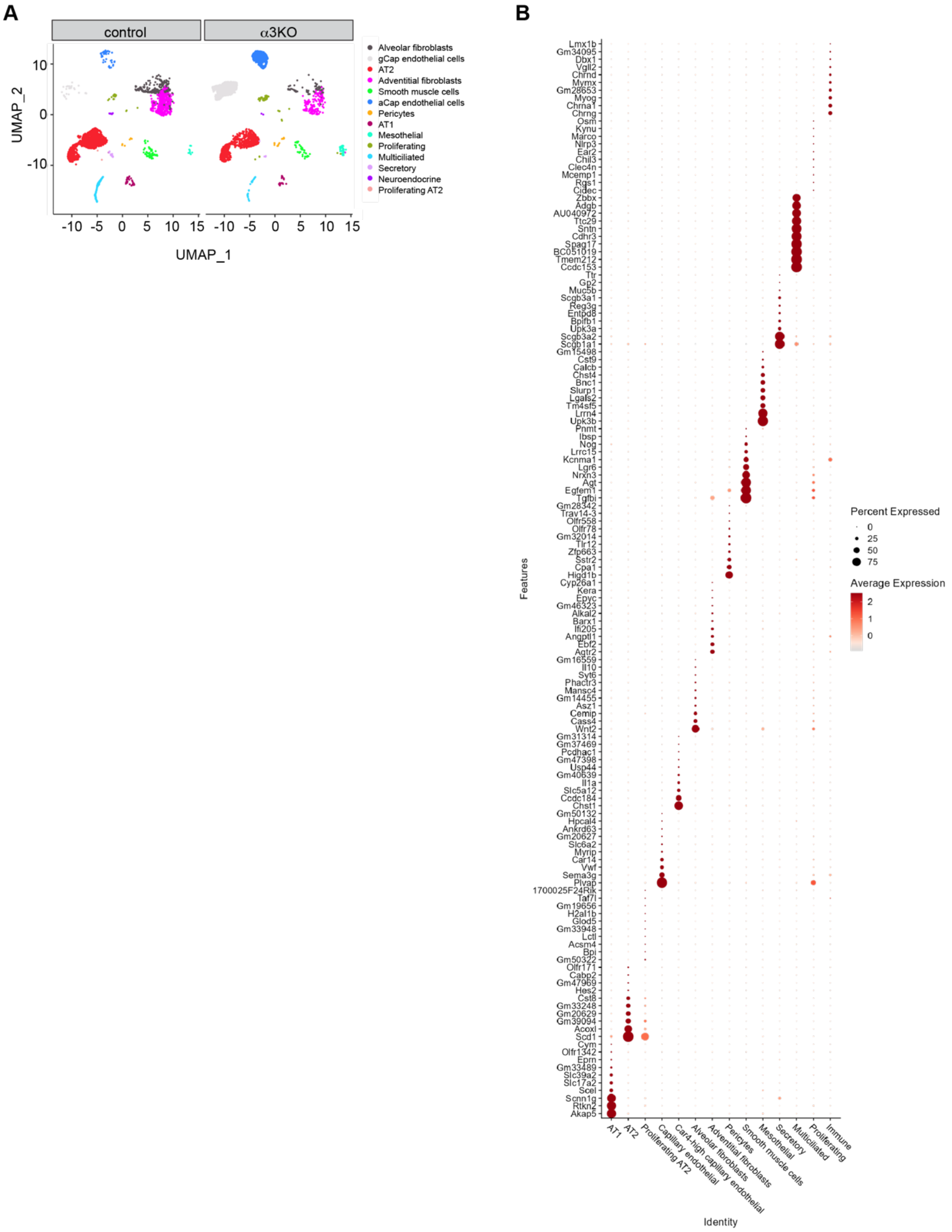


**Supp Fig 5**. (A) UMAP of all cells from three-month-old control and α3KO lungs. (B) Marker gene expression by cluster dot plot, in which higher expression is represented with a darker color and the size of the dot reflects the proportion of cells expressing that gene. n= 4 control and 3 α3KO mice.

Supplemental Figure 6

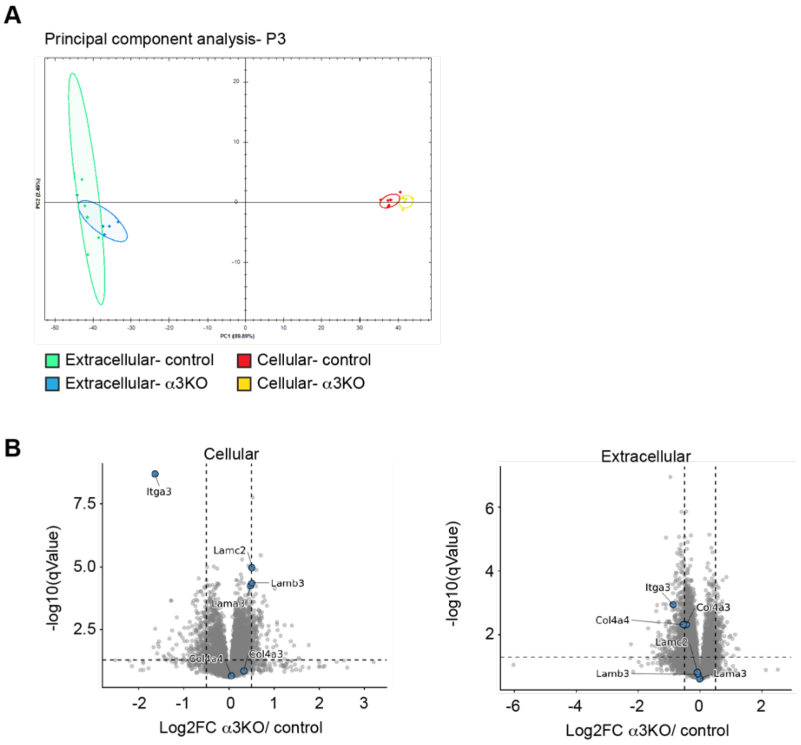


**Supp Fig 6**. (A) Principal component analysis of DIA proteomic data demonstrates P3 extracellular proteins cluster independently from cellular fractions in α3KO and control lungs. The α3KO cellular protein fraction was also distinct from the control cellular fraction at P3. Individual mice represented by dots, n = 4-6 mice per group. (B) Volcano plots of laminin-332 and collagen IV α3α4α5 protein subunits in cellular and extracellular P3 whole lung. Horizontal dashed line corresponds to a q-value of ≤ 0.05 and vertical dashed lines correspond to absolute $\log_2$ FC ≥ 0.6. n = 4-5 mice/ group. Collagen IV α5 was not identified in either genotype.
